# Substrate Channeling *via* a Transient Protein-Protein Complex: The case of D-Glyceraldehyde-3-Phosphate Dehydrogenase and L-Lactate Dehydrogenase

**DOI:** 10.1101/2020.01.22.916023

**Authors:** Željko M. Svedružić, Ivica Odorčić, Christopher H. Chang, Draženka Svedružić

## Abstract

**Background:** D-Glyceraldehyde-3-phosphate dehydrogenase (GAPDH) and L-lactate dehydrogenase (LDH) can form a complex that can regulate the major metabolic pathways, however, the exact mechanism remains unknown. We analyzed a possibility of NADH-channeling from GAPDH-NADH complex to LDH isozymes using enzymes from different cells.

**Results:** Enzyme-kinetics and NADH-binding studies showed that LDH can use GAPDH-NADH complex as a substrate. LDH activity with GAPDH-NADH complex was challenged with anti-LDH antibodies to show that the channeled and the diffusive reactions always take place in parallel. The channeling path is dominant only in assays with limiting free-NADH concertation that mimic cytosolic conditions. Analytical ultracentrifugation showed that the channeling does not require a high affinity complex. Molecular dynamics calculations and protein-protein interaction studies showed that LDH and GAPDH can form a leaky channeling complex only at subsaturating NADH concentrations. The interaction sites are conserved between LDH isozymes from heart and muscle, and between GAPDH molecules from rabbit and yeast cells. Positive electric fields between the NAD(H) binding sites on LDH and GAPDH tetramers, showed that NAD(H)-channeling within the LDH-GAPDH complex, can be an extension of NAD(H)-channeling between the adjacent subunits in each tetramer.

**Conclusions:** In the case of a transient (GAPDH-NADH)-LDH complex, the relative contribution from the channeled and the diffusive paths depends on the overlap between *off*-rates for the transient (GAPDH-NADH)-LDH complex and *off*-rates for the GAPDH-NADH complex. Molecular evolution or metabolic engineering protocols can exploit substrate channeling for metabolic flux control by fine-tuning substrate-binding affinity for the key enzymes in the competing reaction paths.

**Highlights:**

- Substrate channeling molecular mechanism can regulate energy production and aerobic and anaerobic metabolism in cells
- LDH and GAPDH can form a channeling complex only at sub-saturating NADH concentration
- Channeled and diffusive paths always compete and take place in parallel
- NADH channeling does not require a high-affinity complex
- NADH channeling within GAPDH-LDH complex is an extension of NAD(H) channeling within each tetramer
- Allosteric modulation of NADH binding affinity in GAPDH tetramer can regulate NAD(H) channeling

## Introduction

D-glyceraldehyde-3-phosphate dehydrogenase (GAPDH) and L-lactate dehydrogenases (LDH) are two NAD(H) dependent dehydrogenases that participate in glycolytic and gluconeogenic pathways ^1,2^. Glycolysis and gluconeogenesis are the two major metabolic pathways that provide cells with metabolic precursors and a rapid source of energy. Parallel to glycolysis GAPDH participates in microtubule bundling, DNA replication and repair, apoptosis, the export of nuclear RNA, membrane fusion, and phosphotransferase activity ^1,2^. GAPDH is implicated in Huntington’s disease ^3^, prostate cancer, and viral pathogenesis ^1,2^. GAPDH could be a target of nitric oxide ^4^ and a target of drugs developed to treat malaria or Alzheimer’s disease ^3,5^.

GAPDH is well-known to bind actin filaments, microtubule networks and cellular membranes ^1,6-9^. The working hypothesis is that GAPDH can regulate cell physiology by binding at sites with high metabolism and energy consumption ^1,6-8^. GAPDH binding at the sites with high physiological activity can trigger binding of other glycolytic enzymes and lead to a rapid and efficient response to the localized changes in cellular metabolism ^1,6-8^. Dynamic interactions between glycolytic enzymes have been frequently explored in the last 60 years ^9-11^. The underlying molecular mechanism and physiological functions are still poorly understood ^9,10^.

NADH channeling can be one of the key functions in supramolecular organization between glycolytic enzymes. NADH channeling has been explored with limited success in the last 40 years ^8,12-17^. A network of channeling reactions between different NAD(H) dehydrogenases can maintain the separation between different NAD+/NADH pools in cells ^1,6,18-20^. Such separations can be the key mechanism in the regulation of cell physiology and tumor development ^1,6,18-20^. Cytosolic and mitochondrial NAD+/NADH pools are highly un-equilibrated ^21^, and still closely integrated in control of cellular energy production ^20,22^. Cytosolic NAD+/NADH pool can also affect cytosolic NADP+/NADPH pool through dual specificity malate dehydrogenase ^23^. The separation between NADH and NADPH pools can regulate the balance between catabolic and anabolic processes in cells ^20,22^.

GAPDH is the most abundant cytosolic dehydrogenase, that binds the majority of cytosolic NAD(H) molecules ^1,2,6,23,24^. NADH channeling from GAPDH to different NADH-dehydrogenases could regulate the separation between different NAD(H) pools in cells ^20,22^. The changes in NAD+/NADH concentrations have a peculiarly strong effect on GAPDH structure ^1,2,6,23-26^. A decrease in NADH concentration leads to partial dissociation of the GAPDH tetramers ^2,6,27^. NAD(H) binding to different subunits in GAPDH tetramer results in allosteric regulation of GAPDH activity and strong negative cooperativity that can change the NAD(H) binding affinity by thousand-fold ^24-26^. These effects are species specific and known for decades, but their physiological significance is still unknown ^2,6,27^.

LDH-GAPDH complex was observed in cell extracts [14], and with purified proteins at high protein concentrations [15]. Such LDH-GAPDH complex can regulate the major metabolic pathways, however, functional consequences of LDH-GAPDH interaction have not been investigated. In this study, we show that LDH and GAPDH tetramers can form a transient supramolecular complex that can simultaneously support channeled and diffusive reactions as a part of a large glycosome. Such competition between channeled and diffusive paths can fundamentally change our understanding of metabolic regulation and metabolic flux analysis. Most notably, changes in the substrate binding affinity can be a molecular mechanism for fine-tuning of metabolic pathways in enzyme evolution and metabolic engineering protocols.

## Results

### Molecular dynamics simulations of the interaction between rabbit muscle GAPDH and rabbit muscle LDH

LDH-GAPDH complex can be observed in cell extracts [14], and with purified proteins only at high protein concentrations [15]. We used different structure analysis techniques to calculate the structure of (unstable) LDH-GAPDH complex. Distribution of electric potentials on protein surface can give initial insights into the structural elements that can support substrate channeling ^28,29^. We find that NADH binding sites on two adjacent monomers in tetramers of rabbit muscle LDH ^30^ are connected with a 19.2 Å long groove that is dominated by the positive potential (Fig. 1 A-B). The groove is observed only when the active site loops are open (Fig. 1A, residues 98-105) ^30^). In GAPDH tetramers ^31^, the NADH binding sites on two adjacent monomers are enclosed within 18.4 Å nm wide gulf between the monomers (Fig. 1 C). The gulf is entirely dominated by positive electric fields (Fig. 1 C-D). The electric fields in NAD(H) binding sites are projecting dominant positive electric fields in the space around each protein (Fig. 1 B and D). Such positive fields can limit diffusion of negatively charged NAD(H) molecules between the adjacent subunits and facilitate substrate channeling within rmLDH or rmGAPDH tetramers ^28,29^. The positive fields around each protein can visibly decrease when the proteins bind negatively charged NAD(H) molecules (Fig. 1 B-D).

**Figure 1 (A-D).**
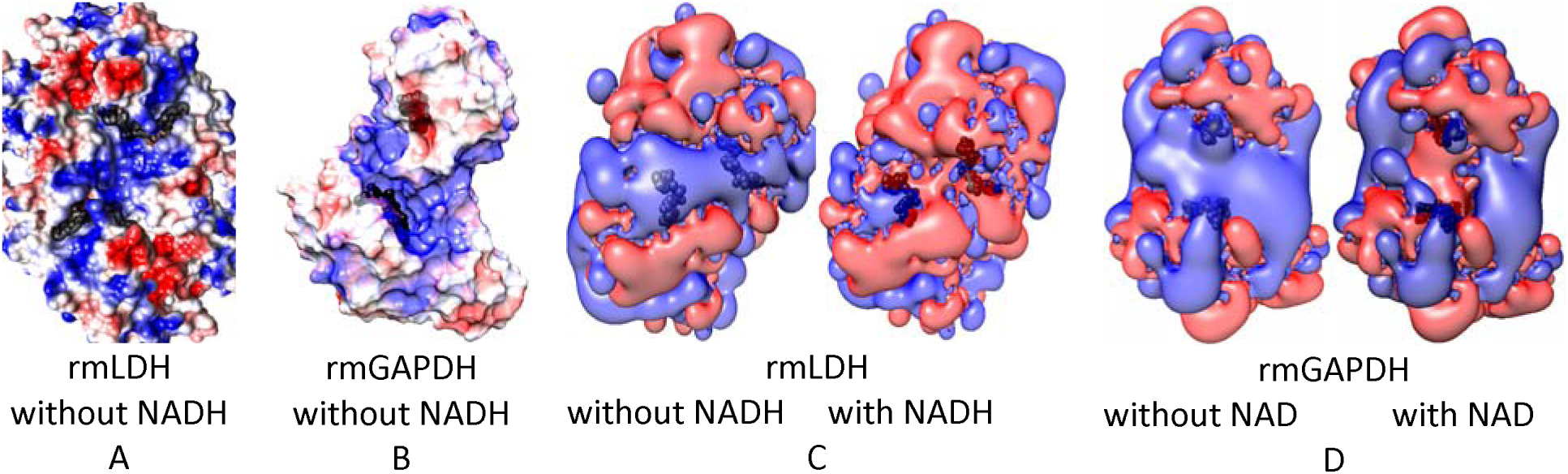
Positive electric fields are connecting the adjacent NADH binding sites in tetramers of rmLDH (PDB: 3H3F, ^30^) and rmGAPDH (PDB:1J0X, ^31^). Adaptive Poisson-Boltzmann Solver (APBS) algorithm was used to calculate electric fields created by the enzymes in an aqueous solution of 150 mM NaCl [35,36,40]. The electric fields were calculated with or without NAD(H) bound to the enzymes. Black CPK models are used to mark the position of NAD(H) binding sites. (**<u>A</u>**<u>-</u>**<u>B</u>**) In both rmLDH and rmGAPDH, the NAD(H) binding sites on the adjacent subunits are connected with a cavity in the protein surface that is dominated by the positive electric fields. The positive fields can channel the negatively charged NAD(H) molecules between the subunits. Red and blue colors indicate potentials from -3.0 to 3.0 k_B_*T/e respectively. (**<u>C</u>**<u>-</u>**<u>D</u>**) The isopotential surfaces show that the space around NAD(H) sites is dominated by the positive potential that can trap the negative NAD(H) molecules on the protein surface. The red and blue colors indicate isopotential surfaces at -0.5 and 0.5 k_B_*T/e respectively. The isopotential surfaces are partially affected by the binding of negatively charged NADH molecules.

Based on the calculated surface potentials, we have presented a possible binding orientation for the rmLDH-rmGAPDH complex in which four NAD(H) binding sites are enclosed in one positive field (Fig. 2A). The number of possible docking orientations is limited by the two main requirements ^28,29^. The distance between the channeling sites should be between 2 to 5 nanometers, and the channeling path should be continuous and at least partially secluded from the surrounding solution.

**Figure 2 (A-E).**
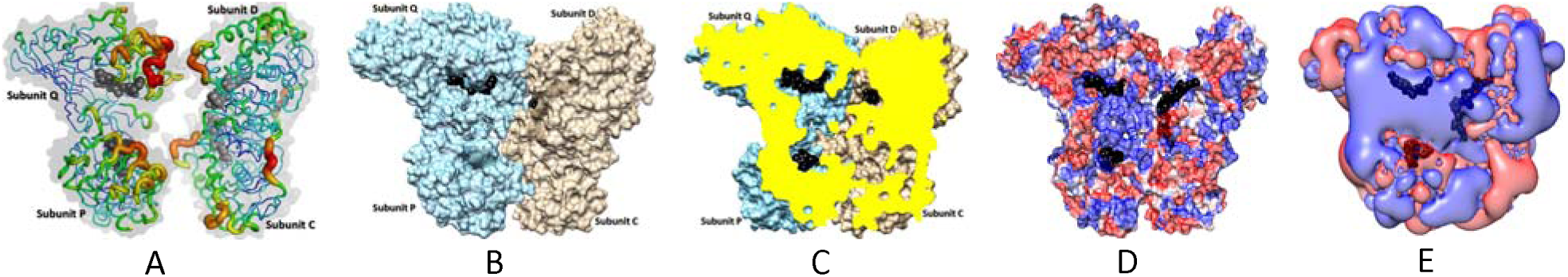
Docking interface between rabbit muscle LDH (PDB:3H3F, ^30^) and rabbit muscle GAPDH (PDB:1J0X, ^31^). The figures show only the interacting subunits to bring in focus the interaction interface. The figures show docking interactions in the absence of NAD(H). Black CPK models were used to mark the position of NAD(H) binding sites. (**<u>A</u>)** The docking surfaces between rmGAPDH (left) and rmLDH (right) consist of flexible loops that have highest values for temperature b-factor. The protein surfaces are shown as transparent gray contours to illustrate relative orientations when the two proteins make their first contacts at the start of docking. (**<u>B-C</u>**) The figures show rmGAPDH-rmLDH complex that was extracted from frame 825 in the supplement movie 1. (**<u>B</u>**) the two proteins are shown in the surface mode to highlight large complementary surfaces (7790 Å^2^) that are not readily apparent from the ribbon models. The surfaces are shown 30% transparent to mark the NAD(H) sites buried under the protein surface. (**<u>C</u>**) the complex is sliced through the plane that is passing through the NAD(H) binding sites to show a central “channeling” cavity that is surrounded by two NAD(H) binding sites from rmGAPDH and one NAD(H) site from rmLDH. **(D-E)** The complex is shown in the same orientation as in panels A to C. (**<u>D</u>**) the complex was sliced through the plane that is passing through the NAD(H) binding sites to show the electric fields that are connecting the sites. (**<u>E</u>**) The red and blue colors indicate isopotential surfaces around the complex at -0.5 and 0.5 k_B_*T/e respectively. The two presentations show that the positive electric fields that can support NAD(H) channeling between the adjacent subunits in rmLDH and rmGAPDH tetramers can merge together to support NAD(H) channeling within the rmLDH-rmGAPDH complex. Thus NAD(H) channeling in the complex is an extension of NAD(H) channeling between the adjacent subunits in rmLDH and rmGAPDH tetramers.

Multiscale MD calculations and molecular docking studies showed that rmLDH and rmGAPDH can form a dynamic complex facing each other with their NAD(H) binding sites (Fig. 2-3, Supp. Fig. 1-8 and 10, and Supp. video 1). The complex breaks apart when the two enzymes are saturated with NAD(H) molecules (Supp. video 1). This is consistent with the earlier experimental studies of interaction between LDH and GAPDH molecules which showed that saturation with NADH leads to complex breakdown ^15^. When the enzymes are saturated with NAD(H) they form some random contacts but ultimately slip apart (Supp. video 1, and Supp. Figs. 8 G-I). Superimposed structures of NADH-free and NADH-bound forms of each enzyme showed that NAD(H)-bound structures cannot support the formation of the complex due to repositioning of the interacting amino acids (Supp. video 2). NADH binding to rmLDH will cause the closing of the active site cleft ^30,32^. NAD(H) binding to GAPDH will cause 7.4 Å contraction between the two subunits that interact with LDH ^31^.

**Figure 3.**
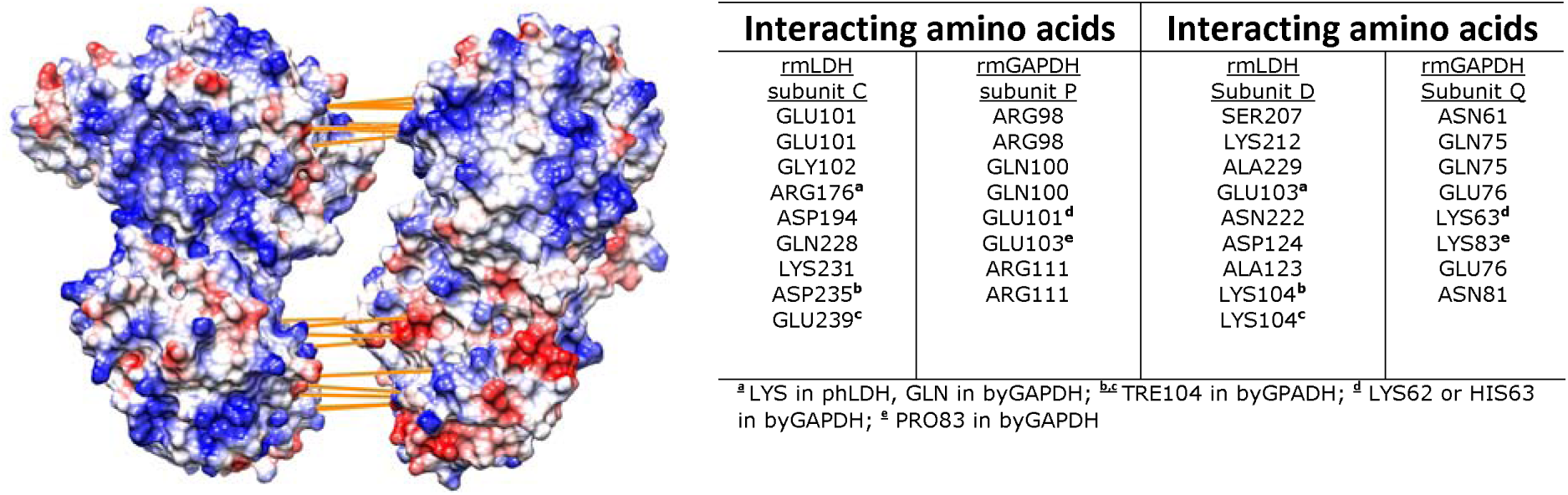
rmLDH and rmGAPDH share complementary surface shape and electric potentials at the interaction sites. The figure shows rmGAPDH:rmLDH complex just as in figures 2 B-E, except that the two proteins were taken apart to expose the binding contacts ^36^. rmLDH and rmGAPDH have large complementary surfaces (7790 Å^2^) and the strongest interactions are focused on two hot spots. In the first spot (740 Å^2^), the most protruded section of NADH binding domain on Q subunit in GAPDH (a.a. 57-74) is wedged between the active site loop (a.a. 98-104) and the active site helix (a.a. 226-245) in D subunit on LDH. The strongest interactions at this site are the three ionic bonds between Glu 76 on GAPDH and ARG 111 on LDH. In the second interaction spot (1210 Å^2^), the catalytic domain on subunit C in LDH binds to the cleft between the catalytic and NADH binding domain on P subunit in GAPDH (a.a. 99-105, a.a. 122-126, a.a. 209-229). Driven by the thermodynamic wobbling between the interacting proteins this interaction site can have as many as 9 binding interactions. The strongest interactions at this site are the three ionic bonds between Arg 176 on GAPDH and Glu 103 on LDH.

Repeated MD simulations showed that the conformational changes around NAD(H) binding sites lead to stochastic differences in the rate of interaction build-up and in variability in the number of binding interactions (Supp. Figs. 7-8 and 10). The structures from the MD frames that show differences in the number of binding interactions have been superimposed to analyze the interaction mechanism. The highest number of binding interactions can be observed when both rmLDH and rmGAPDH tetramers are present in their open conformations that are dominant in NADH-free structures (Fig. 3 and Supp. video 1-2). Most notably stochastic differences in the repeated simulations can be attributed to the random wobbling in the position of the active site loop on rmLDH (a.a. 98-104). When the active site loop is closed the interacting proteins start wobbling around the axis that is perpendicular to the plane of interaction (Supp. video 1 and 2). The delicate nature of the presented LDH-GAPDH interactions can explain why it was so difficult to measure that complex ^14,15^. The LDH-GAPDH tetramers form a symmetric complex that can support association of multiple LDH and GAPDH molecules in a polymer, which can explain the poor solubility of the complex ^14,15^.

When rmLDH and rmGAPDH form a complex, the positive cavities on the surface of each enzyme merge to form a central positive cavity under the protein surface (Fig. 2 C-E). The cavity connects four NAD(H) binding sites with an average separation of 2.9 nm between the adjacent sites (Fig. 2 C-E). Thus, NADH channeling within the rmLDH-rmGAPDH complex can be an extension of NADH channeling between the two adjacent monomers in rmLDH and rmGAPDH tetramers (Fig. 1-2).

### Interaction between LDH and GAPDH molecules from different cells

LDH and GAPDH molecules from different cells have highly conserved structures but differ in the net charge and the NAD(H) binding mechanism ^6,15,24,33^. Both rabbit muscle and porcine heart LDH tetramers can form complex with rmGAPDH [15]. phLDH has 75% sequence identity and 93.1% sequence similarity with rmLDH (5LDH ^34^ and 3H3F ^30^). The two molecules have opposite net charge (pI(rmLDH)=8.1, pI(phLDH)=4.6) [15]). The corresponding structures can overlap with the RMSD value of 1.73 Å. The amino acids that form binding interactions in the rmLDH-rmGAPDH complex appear to be conserved between rabbit muscle LDH and porcine heart LDH (Fig. 3). Repeated MD simulations showed that phLDH-rmGAPDH complex can form in average 2 more binding interactions than the rmLDH-rmGAPDH complex (Supp. Fig. 9 A-C). The most notable difference is the replacement of Gln 228 in rmLDH with Glu 227 in phLDH (Fig 3). The substitution is on a flexible helix that can adapt to different orientations between the interacting proteins (Fig. 3). Porcine and rabbit heart LDH molecules share 99.1% sequence similarity, 95.8 % sequence identity, and 100 % identity at the interaction sites (Fig. 3).

Rabbit muscle GAPDH and baker’s yeast GAPDH have 65.5% sequence identity and 84.8% similarity (PDB: 3PYM, isozyme 2, ^31,35^). The corresponding structures can overlap with the RMSD value of 0.556 Å. byGAPDH and rmGAPDH have very different NADH binding affinities and opposite net charge (pI(byGAPDH)=6.5, pI(rmGAPDH)=8.2) ^31,35^. Repeated MD calculations showed that byGAPDH-rmLDH complex can form in average 8 ± 2 binding interactions, i.e., two interactions less than rmGAPDH:rmLDH complex (Fig. 3, Supp. Fig. 9 D-F). Most notably, Lys 83 and Lys 104 in rmGAPDH are replaced with Pro83 and Thr105 in byGAPDH (Fig. 3).

### Enzyme buffering tests with different LDH molecules as enzyme acceptor and GAPDH molecules as NADH donor enzyme

We have designed enzyme assays that can measure NADH channeling from GAPDH-NADH complex to different LDH isozymes (Supp. Figs. 11-14.) The measurements were designed to mimic LDH activity in the cytosol, where the majority of NAD(H) molecules are bound to GAPDH, the most abundant dehydrogenase in cells with the highest NADH binding affinity ^6,23^. The enzyme assays with two enzymes that share a common substrate can be challenging to design and interpret ^12,13^, especially in the case of transient LDH-GAPDH interaction (Supp. Figs. 11. ^14,15^) Thus, we used extensive numerical simulations ^37^ in preparation for optimal assay design and data interpretation (Supp. Figs. 11 to 14). The simulations showed that the free-diffusion and the channeling paths always take place in parallel. Thus, we measured LDH activity using two approaches that can modulate the relative contributions from the two paths: increasing GAPDH concentration with fixed NADH concentration and decreasing NADH concentration with fixed GAPDH concentration. In the case of no channeling the measured activity will be equal to the calculated LDH activity on free NADH substrate (eqn. 1-2, supp. Fig. 11). The expected free-diffusion activity can be calculated using Michaelis-Menten constants for LDH activity with its NADH substrate (Table 2) and dissociation constant Kd for GAPDH-NADH complex (Tables 1-2). In the case of channeling, the measured LDH activity will be higher than the calculated activity on free NADH substrate (Supp. Fig. 11). We found with four different enzyme pairs that LDH activity in the presence of a large excess of GAPDH is significantly higher than the expected activity on free NADH substrate (Figs. 4 and 5).

**Table 1.**
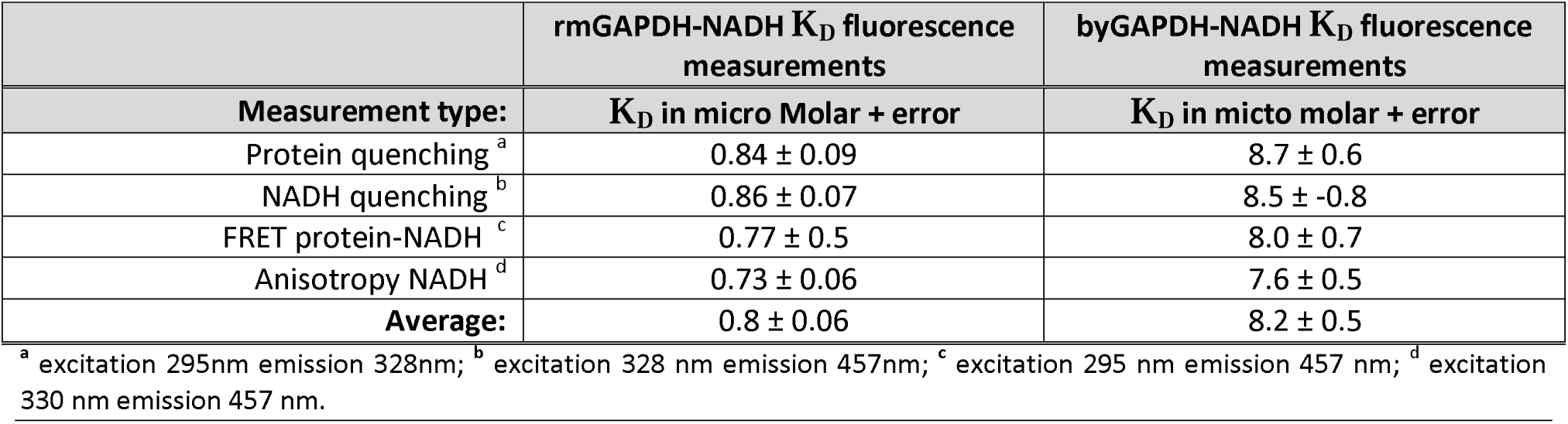
Four different types of fluorescence measurements were used to measure NADH binding affinity for rmGAPDH and byGAPDH. The K_D_ constants are shown in terms of NADH binding sites on each tetramer.

| | rmGAPDH-NADH $K_D$ fluorescence measurements | byGAPDH-NADH $K_D$ fluorescence measurements |
| --- | --- | --- |
| Measurement type: | $K_D$ in micro Molar + error | $K_D$ in micro molar + error |
| Protein quenching <sup>a</sup> | 0.84 ± 0.09 | 8.7 ± 0.6 |
| NADH quenching <sup>b</sup> | 0.86 ± 0.07 | 8.5 ± -0.8 |
| FRET protein-NADH <sup>c</sup> | 0.77 ± 0.5 | 8.0 ± 0.7 |
| Anisotropy NADH <sup>d</sup> | 0.73 ± 0.06 | 7.6 ± 0.5 |
| **Average:** | 0.8 ± 0.06 | 8.2 ± 0.5 |

<sup>a</sup> excitation 295nm emission 328nm; <sup>b</sup> excitation 328 nm emission 457nm; <sup>c</sup> excitation 295 nm emission 457 nm; <sup>d</sup> excitation 330 nm emission 457 nm.

**Table 2.**
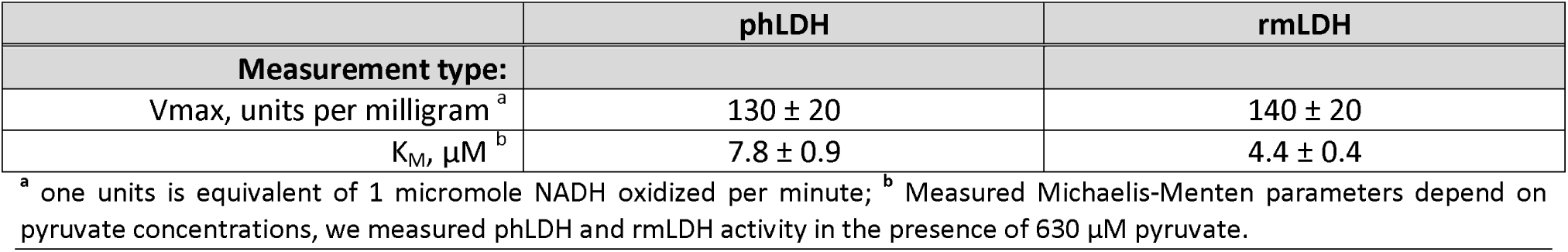
Michaelis-Menten constants for phLDH and rmLDH with free NADH as substrate.

|  | phLDH | rmLDH |
| --- | --- | --- |
| Measurement type: |  |  |
| Vmax, units per milligram <sup>a</sup> | 130 ± 20 | 140 ± 20 |
| $K_M$, μM <sup>b</sup> | 7.8 ± 0.9 | 4.4 ± 0.4 |

<sup>a</sup> one units is equivalent of 1 micromole NADH oxidized per minute; <sup>b</sup> Measured Michaelis-Menten parameters depend on pyruvate concentrations, we measured phLDH and rmLDH activity in the presence of 630 μM pyruvate.

**Figure 4 (A-B).**
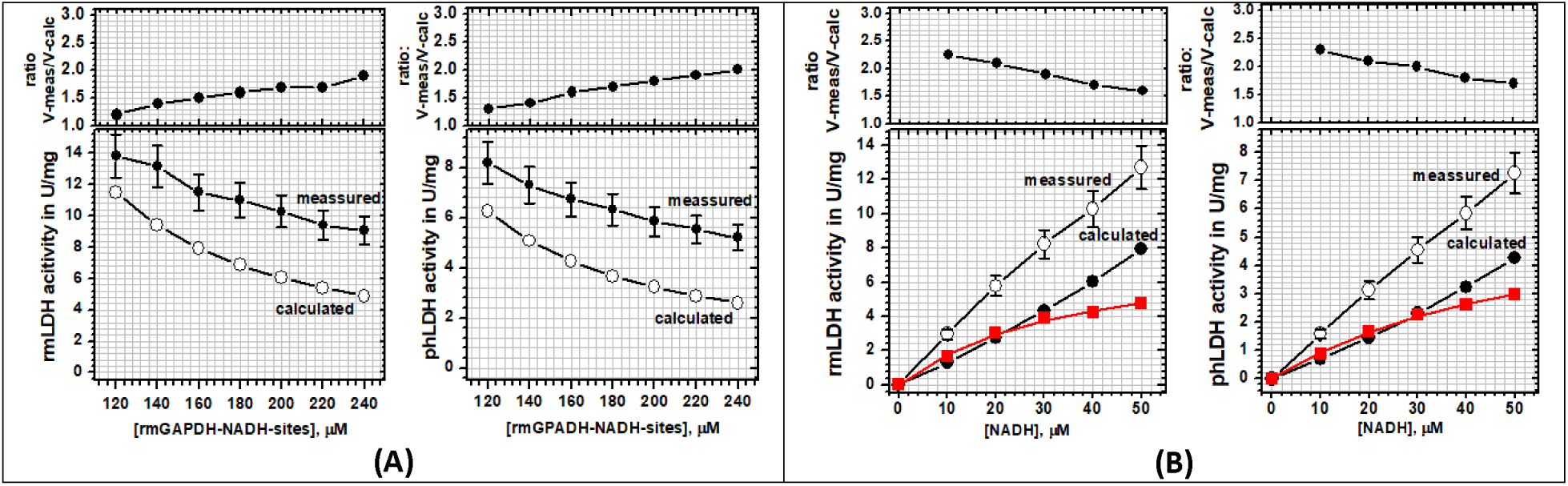
The activity of rabbit muscle LDH and porcine heart LDH was measured in the presence of a large excess of rabbit muscle GAPDH. **(<u>A</u>)** steady-state activities of rmLDH (10 nM) or phLDH (17 nM) were measured at fixed NADH concentration (40 µM) in the presence of increasing concentration of rmGAPDH (100 to 240 μM of NADH binding sites). Increase in rmGAPDH concentration leads to the disproportional decrease in the measured and the calculated free-diffusion activities (lower panels) which results in the increase in the ratio between the two activities (upper panels, and equations 2-3). These results indicate that LDH molecules can use GAPDH-NADH complex as a substrate in addition to free NADH, i.e. NADH channeling (Supp. Fig. 11). **(<u>B</u>)** steady-state activities of rmLDH (10 nM) or phLDH (17 nM) were measured in the presence of decreasing NADH concentration with rmGAPDH fixed at 200 μM (NADH binding sites). The decrease in NADH concentration leads to the disproportional decrease in the measured and the calculated free-diffusion activities (lower panels), what results in increase in the ratio between the two activities (upper panels, and equations 2-3). The red curve represents the Michaelis-Menten profile for LDH activity with the rmGAPDH-NADH complex as the substrate, which was calculated by subtracting the calculated free-diffusion profile from the measured profiles (supp. Fig 14). The calculated apparent K_M_ constant for rmLDH is 35 ± 5.5 μM and 59 ± 6 μM for phLDH (Table 3). Thus, the observed K_M_ constants are a result of competition between the channeled and free-diffusion paths (Supp. Fig. 11).

**Figure 5 (A-B).**
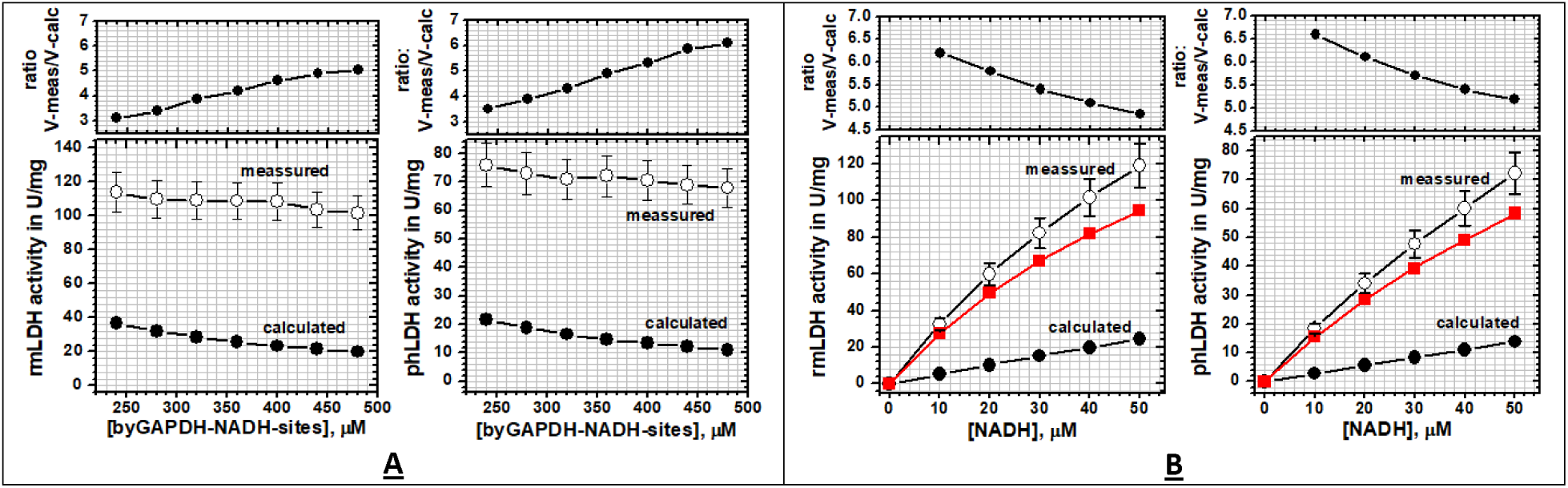
The activity of rabbit muscle LDH and porcine heart LDH was measured in the presence of a large excess of baker’s yeast GAPDH. **(<u>A</u>)** steady-state activities of rmLDH (10 nM) or phLDH (17 nM) were measured at fixed NADH concentration (40 µM) in the presence of increasing concentration of byGAPDH (240 to 480 μM in terms of NADH binding sites). Increase in byGAPDH concentration leads to the disproportional decrease in measured and calculated free-diffusion activities (lower panels) which results in increase in the ratio between the two activities (upper panels, and equations 2-3). These results indicate NADH channeling from byGAPDH-NADH complex to rmLDH or phLDH (Supp. Fig. 11). **(<u>B</u>)** steady-state activities of rmLDH (10 nM) or phLDH (17 nM) were measured in the presence of decreasing NADH concentration with byGAPDH fixed at 480 μM (NADH binding sites). The decrease in NADH concentration leads to the disproportional decrease in the measured and the calculated free-diffusion activities (lower panels), what results in the increase in the ratio between the two activities (upper panels, and equations 2-3). The red curve represents the Michaelis-Menten profile for LDH activity with the byGAPDH-NADH complex as the substrate, which was calculated by subtracting the calculated free-diffusion profile from the measured profiles (supp. Fig 14). The calculated apparent K_M_ constant for rmLDH is 78 ± 3 μM and 116 ± 10 μM for phLDH (Table 3). Thus, the observed K_M_ constants are a result of competition between the channeling and free-diffusion paths.

In the first set of experiments, we compared rmLDH and phLDH as enzyme acceptors with rmGAPDH as a donor enzyme (Fig 4 A-B). LDH activities were measured with fixed NADH concentration (40 μM) and variable rmGAPDH concentrations (100 to 240 μM of NADH binding sites or 3.5-8.5 mg/ml protein) (Fig. 4 A). In these conditions, more than 99% of all NADH molecules are bound to rmGAPDH (supplement Table 2). The free NADH concentration is at least 11-fold lower than the related K_M_ constants for each LDH molecule (supplement Table 2). Increase in rmGAPDH concentration leads to an increase in the ratio between measured and calculated activity from 1.2 to 1.85 fold (eqn. 3 in methods). In the same experiments with phLDH, increase in rmGAPDH leads to an increase in the ratio between the measured and calculated activity from 1.3 to 2.0 fold (eqn. 3 in methods). The experiments with fixed rmGAPDH concentration showed that NADH decrease from 40 to 10 μM leads to increase in the ratio between measured and calculated activity from 1.65 to 2.25 fold for rmLDH and 1.7 to 2.28 fold for phLDH (Fig. 4B and supplement Table 2).

In the second set of experiments, we have replaced rmGAPDH as enzyme donor with byGAPDH (Fig. 5 A-B). The activities or rmLDH and phLDH were measured at fixed NADH concentration (40 μM) and variable concentrations of byGAPDH (240 to 480 μM of NADH binding sites or 8.5-17.6 mg/ml protein). In those conditions, more than 96% of NADH molecules are bound to byGAPDH (supplement Table 2B). The concentration of free NADH is at least five-fold lower than the respective Km constants for rmLDH or phLDH (supplement Table 2B). In experiments with rmLDH, increase in byGAPDH concentration leads to an increase in the ratio between measured and calculated activity from 3.5 to 5.8 fold (Fig 5A and supplement Table 2B). In the same experiments with phLDH, increase in byGAPDH concentration leads to an increase in the ratio between measured and calculated activity 3.5 to 6.1 fold (Fig 5A and Supp. Table 2B). The experiments with fixed byGAPDH concentration showed that NADH decrease from 40 to 10 μM leads to increase in the ratio between measured and calculated activity from 4.85 to 6.2 fold for rmLDH and 5.2 to 6.6 fold for phLDH (Fig. 5B supplement Table 2).

In conclusion, consistent with the molecular dynamic studies, all four experiments showed that the channeled and the diffusive reactions always take place in parallel. The channeling can be measured only at extremely high protein concentrations which mimic cytosolic conditions, that favor the formation of GAPDH-NADH complex at the expense of free NADH (Figs. 4-5) ^1,2,6,23,24^. These results are consistent with the earlier experiments ^14,15^ and with the numerical simulations (Supp. Figs. 11-14). We find that the highest GAPDH concentrations are limited by the enzyme solubility, while the lowest NADH concentrations were limited by assay sensitivity (supplement Table 2). In all four experiments the conditions that favor formation of GAPDH-NADH complex lead to decrease in measured LDH activity, about 38% decrease with rmGAPDH and about 10% decrease with byGAPDH (Figs. 4 and 5). The decrease can be attributed to the shift from the faster turnover in free-diffusion reaction to the slower turnover in channeling reaction (Supp. Figs. 11 to 14). The turnover rates in channeling reaction are always slower than the turnover rates for the diffusive reaction due to one extra step: NADH dissociation from GAPDH within (GAPDH-NADH)-LDH complex (Supp. Figs. 12-13). In all four measurements, the apparent K_M_ values for LDH molecules with GAPDH-NADH substrate are higher than the K_M_ values for free NADH (Table 2 vs. 3, and Figs. 4B and 5B). Thus, in the case of channeling the extent of saturation of LDH activity with the NADH substrate is a result of competition between the channeled and diffusive reaction paths (Supp. Fig. 14).

### Control Enzyme Buffering Experiments: competition assays with anti-phLDH Antibodies

In a negative control experiment, the enzyme buffering experiments were repeated in the presence of an excess of polyclonal anti-LDH antibodies (Fig. 6). The large antiLDH molecules have little effect on the binding of small NADH molecules to phLDH. On the other hand, approximately 45% inhibition was observed in the presence of rmGAPDH (240 μM NADH binding sites or 8.5 mg/ml protein). Similarly, about 80% inhibition was observed in the presence of byGAPDH (450 μM NADH binding sites or 15.9 mg/ml). The anti-LDH antibodies have a much higher binding affinity for LDH molecules than the GAPDH molecules, and thus the anti-LDH antibodies can readily interfere with the formation of different LDH-GAPDH complexes (Fig. 6).

**Figure 6.**
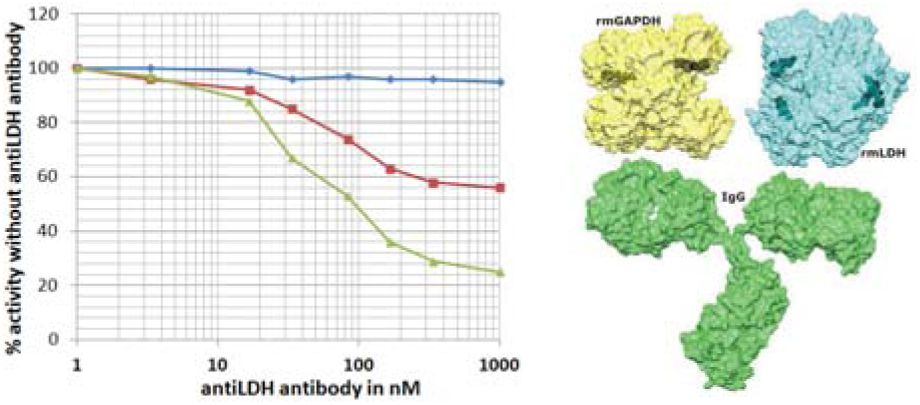
phLDH activity was measured in the presence of increasing concentration of polyclonal anti-phLDH antibodies (0 to 1000 nM or 0.15 to 156 μg/ml). phLDH activity (16 nM) was measured with free NADH (blue line), or in the presence of 240 μM rmGAPDH (red line) or 450 μM byGAPDH (green line). In all cases total NADH concentration was 40 μM, and the GAPDH concentrations are expressed in terms of NADH binding sites in each tetramer. Bulky anti-LDH antibodies can slightly inhibit activity of phLDH with small NADH molecules and produce a significantly higher inhibition with the large GAPDH-NADH complex. The relative size of each molecule shows that bulky IgG molecules (PDB:1IGT) can interfere with LDH-GAPDH interaction for all epitopes that are close to the NADH binding sites.

At the highest antibody concentration, the measured phLDH activity in the presence of GAPDH molecules is close to calculated activity for the reaction that depends on free diffusion (methods eqns. 1-2). A good agreement between the calculated and the measured activity indicates that the K_D_ constant for GAPDH-NADH complex and Michaelis-Menten parameters for LDH activity were measured with high accuracy (Tables 1 and 2).

These results support our proposals that that the channeled and the diffusive reactions always take place in parallel.

### Analysis of Interaction between phLDH and byGAPDH by Analytical Ultracentrifugation

Our earlier studies of the interaction between phLDH and rmGAPDH showed that AUC studies can be a very sensitive method for detection of interaction between LDH and GAPDH molecules ^15^. The same approach is now used for the analysis of the interaction between phLDH and byGAPDH, the two enzymes that show channeling (Figs. 5-6). The sedimentation constant for phLDH is s^0^_20,w_=7.5 ± 0.2 S, while apparent molecular mass is *Mr*=143,6 ± 0.8 kDa. The calculated *Mr* is within 3% of the calculated *Mr* based on amino acid sequence (146,5 kDa). For byGAPADH the sedimentation constant s^0^_20,w_ =7.65 ± 0.2 S, while apparent molecular mass is 142.1 ± 0.7 kDa what is to within 1.4 % of the *Mr* value that can be calculated based on amino acid sequence (143 kDa).

The sedimentation profiles for phLDH and rmGAPDH have such high similarity that that two proteins mixture can give a good fit to one component model, with the best fit values s^0^_20,w_ =7.6 ± 0.4 S, *Mr*=143,6 ± 0.9 kDa (Fig. 7A). The good fit to one-component model shows that there is no detectable complex between phLDH-byGAPDH in the two-protein mixture. A good fit to one component model can be also used to estimate the sensitivity of our AUC experiments to detect traces of association. The calculated *s* and *Mr* values for a single component model can be used in Claverie simulations in SEDFIT program to simulate sedimentation profile for self-dimerization model assuming 10% association ^38^. In our experiment, each protein in the mix is present in total concentration of 6 μM, thus if there is 10% association the complex concentration will be 0.6 μM, the concentration of free protein is 5.4 μM and the corresponding dissociation constant K_D_ is 48.6 μM (i.e. K_D_ =(5.4)*(5.4)/0.6). The measured and the simulated profiles show detectable differences (Fig. 7). We conclude that there is no interaction between phLDH and byGAPDH with dissociation constant lower than 48.6 μM.

**Figure 7 (A-B).**
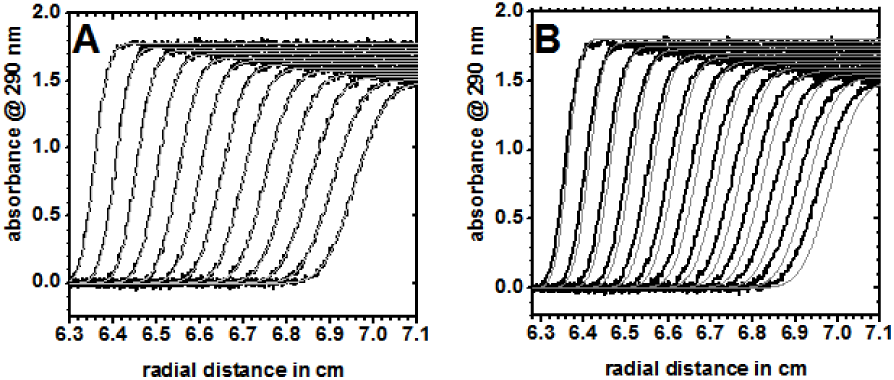
Sedimentation velocity experiments for detection of interaction between phLDH and byGAPDH. The sedimentation profiles for a mixture of phLDH and byGAPDH were measured by following scans at 280 nm. The two-enzyme mixture was prepared using 6 μM solution in 50 mM Tris/HCl, pH=7.2, 1 mM EDTA, 0.5 mM DTT. In both panels measured sedimentation profiles are shown in black, and best-fit profiles are shown in light gray lines. **(<u>A</u>)** the panel shows overlap between sedimentation profiles and the best-fit profiles assuming one component with sedimentation constant s^0^_20,w_=7.6 ± 0.4 S, *Mr*=143,6 ± 0.9 kDa. **(<u>B</u>)** the panel shows measured sedimentation profiles for the two proteins mixture just as panel A, except that the gray lines show Claverie simulation profiles for a single component with 10% self-association. The observed differences between measured sedimentation profiles and the calculated profiles indicate that there is no detectable interaction between phLDH and byGAPDH with the K_D_ constant lower than 48.6 μM.

Presented results (Fig. 4-7) and the earlier LDH-GAPDH interaction studies ^14,15^ indicate that NADH channeling does not require a high affinity complex between GAPDH and LDH. These results indicate that the channeling is taking place within a transient (GAPDH-NADH)-LDH complex (supp. Fig. 11 and ^39^).

## Discussion and Conclusions

### The molecular mechanism and the regulation of NADH channeling from GAPDH-NADH complex

The observed differences between rmGAPDH and byGAPDH can give us insights into the channeling mechanism and the enzyme buffering experiments (Figs. 4-5, supp. Table 2 and 3). rmGAPDH and byGAPDH have very similar structures but differ in their NAD(H) binding mechanism ^6,25,35,40^. Briefly, rmGAPDH (PDB: 1J0X) and byGAPDH (PDB: 3PYM, isozyme 1) have 65.5% sequence identity and 84.8% similarity. The corresponding structures can overlap with RMSD value of 0.556 Å. The two enzymes have such high structural similarities that they can even form heterotetramers ^6,25,35,40^. The heterotetramers readily fall apart after saturation with NAD(H) due to different conformational changes caused by the NADH binding to yeast and rabbit subunits ^6,25,35,40^. rmGAPDH shows strong negative cooperativity in NADH binding to different subunits in the tetramer ^6,40^. byGAPDH does not show negative cooperativity in NADH binding ^35^ and its binding affinity is 10 times weaker than the NADH binding affinity for rmGAPDH (Table 1).

**Table 3.** Apparent Michaelis-Menten constants for phLDH and rmLDH with rmGAPDH-NADH or byGAPDH-NADH complex as substrate:

|  | phLDH | rmLDH |
| --- | --- | --- |
| Measurement type: | rmGAPDH-NADH | rmGAPDH-NADH |
| Vmax, units per milligram <sup>a</sup> | 6.5 ± 0.4 | 8.2 ± 0.7 |
| $K_M$, μM <sup>b</sup> | 59.6 ± 6 | 35 ± 7 |
|  | phLDH | rmLDH |
| Measurement type: | byGAPDH-NADH | byGAPDH-NADH |
| Vmax, units per milligram <sup>a</sup> | 192 ± 10 | 241 ± 3 |
| $K_M$, μM <sup>b</sup> | 116 ± 8 | 78 ± 2 |

<sup>a</sup> one units is equivalent of 1 micromole NADH oxidized per minute; <sup>b</sup> Measured Michaelis-Menten parameters depend on pyruvate concentrations, we measured phLDH and rmLDH activity in the presence of 630 μM pyruvate.

Numerical analysis showed that in the case of transient LDH-(NADH-GAPDH) interaction weaker NADH binding to GAPDH can favor channeling (supp. Figs. 11-13). The studies with increasing *off*-rates showed that in case of transient protein-protein interactions the channeling depends on an overlap in timing of two events: formation of LDH-(NADH-GAPDH) complex and NADH dissociation from GAPDH-NADH complex (supp. Figs. 11-13). The higher K_D_ constant for the byGAPDH-NADH complex is in large part a result of higher *off*-rates for GAPDH-NADH complex ^25,40-44^. Numerical analysis showed that the increase in *off*-rates for GAPDH-NADH complex can qualitatively reproduce experimentally observed differences between rmGAPDH and byGAPDH molecules (supp. Figs. 12-13).

The correlation between NADH channeling and dissociation constant for GAPDH-NADH complex indicates that the negative cooperativity in NADH binding to different subunits in GAPDH tetramer could regulate the NADH channeling ^1,2,40^. Allosteric regulation of substrate channeling has been reported in some of the earlier studies ^11,29^. The negative cooperativity in NADH binding to different subunits in GAPDH molecules is in essence a type of allosteric regulation ^25,42,45^. The negative cooperativity can affect NADH binding affinity to different subunits in GAPDH tetramer by about 1000-fold ^43,44^, from 10 nM to 30 μM. The 1000-fold difference in the binding affinity can be in large part a result of differences in the *off*-rates for the GAPDH-NADH complex ^1,40,41^. In other words, we hypothesize that we can measure no-channeling, low-channeling, and high-channeling between rmGAPDH and LDH molecules if we can design experiments that can selectively measure differences in channeling between each of the four subunits in rmGAPDH tetramers. Such experiments will be significantly more demanding than the presented experiments (Supp. Fig. 11), however such efforts can be very significant. The negative cooperativity in NADH binding to different subunits on rmGAPDH was reported almost 60 years ago, yet to this day the physiological significance of such mechanism is not understood ^6,43,44^. Differences in NAD(H) binding cooperativity can be observed with GAPDH molecules from different tissues ^6,24,42^. Different types of binding cooperativity can be also observed between NAD and NADH substrates ^6,24,42^. Thus, studies of changes in substrate channeling caused by the changes in NAD(H) binding cooperativity can give crucial insights in the regulation of anabolic and catabolic processes in cells ^6,8,11^.

### Transient Protein-Protein interactions and NADH channeling in cells

Presented results (Fig. 4-7) and the earlier protein-protein interaction studies ^14,15^ indicate that NADH channeling is taking place within a transient (GAPDH-NADH)-LDH complex (supp. Fig. 11 and ^39^). Transient interactions could be a physiological necessity for the key metabolic enzymes such as GAPDH which has to rapidly interact with lots of different proteins to accommodate to rapid changes in cell physiology ^6,11,23,39^. Transient interactions can be very difficult to describe in studies of protein-protein interaction (Fig. 7 and ^39^).

The transient GAPDH-LDH complex that forms in experiments with purified enzymes can be much more durable in concentrated protein solution in cytosol or in cell extracts ^14,15,46^. The high protein concentrations can produce *excluded volume effects* that can increase the interaction energy (Δ*G*) by favoring protein-protein interactions that can decrease the number of water molecules trapped in hydration shells ^14,15,46^. It is also necessary to notice that in cytosol LDH-GAPDH complex can be a part of much larger glycolytic metabolon ^8-10^. Such a complex can provide shared molecular scaffolds and molecular crowding effects that can favor collision between the two molecules (*on*-rates) and decrease the complex breakdown rates (*off-*rates). Both of those two processes can provide additional stability to LDH-GAPDH complex.

The presented LDH-GAPDH complex indicates that LDH and GAPDH tetramers can simultaneously participate in channeling reaction, in diffusive reaction, and in interaction with other enzymes (supp. video 1 and Fig. 2 B-E). In the presented GAPDH-LDH complex only one of the four subunits in each tetramer is participating in the interaction with its NAD(H) site (Fig. 2 C-E). Only NADH binding sites on D subunit on LDH is directly facing the Q subunit on GAPDH (Fig. 2). The NADH binding site on P subunit of GAPDH is open to bind free NAD(H) from the solution, but it is also enclosed in a dominant positive electric field that can channel NAD(H) molecules electrostatically to D subunit on LDH ^28^ (Fig. 2). A similar situation is observed with the C subunit on LDH (Fig. 2 B-E). In other words, free NADH can bind to either Q or P subunits on GAPDH and then get transferred by substrate channeling to D or even C subunits on LDH (Fig. 2 B-C). Subunits A and B on LDH and O and R on GAPDH are open for interaction with other enzymes and could not participate in NADH channeling in the presented GAPDH-LDH complex (supp. video 1 and Fig. 2 B-E). The need to fulfill the multiple functions simultaneously can explain why LDH and GAPDH molecules exist in cells as tetramers. APBS analysis showed that NADH channeling between GAPDH and LDH appears to be an extension of NAD/NADH channeling between the adjacent subunits in GAPDH or LDH tetramers (Figs. 1-2).

Substrate channeling is most frequently described as a physiological mechanism that leads to efficient utilization of reactive metabolites ^23,29,39^. The concentration of glycolytic enzymes in cells is much higher than the concentration of their substrates ^23^, what indicates that in glycolysis the regulation of enzyme activity can be more important than efficient substrate utilization ^11,47^. Our results indicate that substrate channeling can regulate metabolism by at least two different mechanisms: *i*) allosteric regulation of differences in enzyme-substrate *off*-rates between different substrates or isozymes (supp. Fig. 12-13), *ii*) the changes in the substrate K_M_ caused by the competition between LDH-NADH, LDH-GAPDH and LDH-(GAPDH-NADH) interactions (Figs. 4B-5B, and supp. Figs. 11-14).

Baker’s yeast cells do not have cytosolic LDH like mammalian cells ^48^. Nevertheless, metabolic engineering studies showed that mammalian LDH can readily integrate into glycolic pathways in baker’s yeast cells ^48^. NADH channeling between byGAPDH and mammalian LDH isozymes can be attributed to highly conserved structures in GAPDH family of enzymes ^6^. The conserved structure could be a result of evolutionary pressure to conserve the related interactome ^6,11,27^.

## Materials and Methods

### Apo-enzyme preparations

(NH_4_)_2_SO_4_ suspensions of purified rmGAPDH and byGAPDH have been prepared in our laboratory using established protocols ^35,40^ or purchased from Sigma Chem. (St. Louis MO)

### Molecular dynamics calculations

All-atom molecular dynamics calculations used the GROMACS 5.1.4 program package ^49^ as we have previously described ^50,51^. Briefly, NAD(H) molecules from PDB files were processed using ACPYPE10, an interface for Antechamber (part of AmberTools11) that can generate topology types for GAFF force fields ^52^. Protein PDB coordinates were processed with pdb2gmx using Amber99SB force field. A cubic solvent box (30 nanometers) was used with TIP3P model for water molecules plus 150 mM NaCl and additional ions that were required for neutralization. The prepared system was minimized using a combination of steepest descent and conjugate gradient algorithms. When the most stable state was achieved the temperature was introduced and the system was equilibrated to 310 K (NVT equilibration, V-rescale). The pressure was equilibrated to 1 atm (NPT equilibration, Parrinello-Rahman). No restraints were used for the protein or the ligand when the system was minimized, but in NPT and NVT equilibration protein and ligand were restrained to prevent the break-up of the complex prior to production runs.

Typical simulations had about 1.5 million atoms, 50 to 150 million steps, with 2 fs timesteps. Large simulation boxes (30 nanometers) were used to avoid attractive or repulsive forces created by the periodic boundary conditions. The large boxes can also provide enough space for the two tetramers to dissociate (Supp. Video 1). Different initial simulation set-ups were used to explore the how representative simulations were of the expected pseudo-equilibrium distribution. The simulations have been repeated with active site loop on LDH in open and closed position, with different initial distances between the interacting proteins, and with different rotations between the LDH and GAPDH tetramers in the interaction plane. Following the simulations, the number of binding interactions was calculated using built-in GROMAC functions. The simulations that showed the highest number of binding interactions have been repeated multiple times to assess variability in the number of binding interactions.

### Adaptive Poisson-Boltzmann Solver (APBS) calculations

All electric fields maps were calculated using **<u>A</u>**daptive **<u>P</u>**oisson-**<u>B</u>**oltzmann **<u>S</u>**olver (APBS) approach ^53^. PDB formats were converted to PQR format using PDB2PQR and PEOEPB force filed with PROPKA set at pH=7.2. NAD(H) molecules from PDB files were protonated using GAFF fields. Potential maps were calculated in aqueous 150 mM NaCl solutions using single Debye-Hückel boundary conditions.

### Fluorescence measurements of NADH binding affinity for rmGAPDH and byGAPDH

Protein fluorescence measurements had excitation 290nm and emission at 335nm. NADH fluorescence had excitation at 340 nm and emission at 460 nm [18, 19, 40, 41, 54]. Protein-NADH FRET measurements had extraction 290 nm and emission at 460 nm.

### Enzyme buffering measurements

In all enzyme buffering experiments LDH activity was measured by following NADH oxidation in the presence of 630 μM pyruvate which results in a decrease in NADH absorbance at 340 nm. The changes in absorbance were measured with Shimadzu UV-VIS 160 Spectrophotometer or with HP 8452A Diode Array Spectrophotometer. The molar absorptivity for NADH is 6.22×103 M-1cm-1. All activity measurements were routinely reproducible to with a precession greater than 3%.

The assay mix was prepared in a microcuvette in a total volume of 80 μL and equilibrated to the temperature at 25° C. The assay buffer was 50 mM Tris/HCl pH=7.4, 2 mM EDTA-Na, 1 mM DTT, and 1mg/ml BSA. Low ionic strength buffer was chosen to favor detection of electrostatic channeling [28, 29]. First, we filled the cuvette with GAPDH solution in the required concentration and incubated the cell in the holder for several minutes to accommodate to the required temperature and to test the stability of the measured absorbance. Second, NADH was added to the cuvette to confirm that GAPDH solution does not have any intrinsic NADH oxidize activity. Third, pyruvate was added in the concentration of 630 μM next to confirm that there is no measurable LDH activity in GAPDH solution. Finally, the reaction was started by adding 10 to 20 nM of LDH.

In assays with NADH concentration between 40 to 50 μM the initial steady-state rates were measured by following the initial linear decrease in absorbance. In the assays with NADH concentration below 30 μM, the rates were calculated by using exponential equations since NADH concentrations were too low to capture the initial linear decrease in NADH oxidation. byGAPDH solutions were stable even at the highest enzyme concentration tested at 16mg/ml. rmGAPDH gradually precipitates at the concentration above 8.4 mg/ml, which leads to scattering and detectable increase in absorbance. Thus, the LDH concentrations to maximize the ratio between the changes in absorbance caused by NADH oxidation and by scattering, so that the scattering had to be subtracted as a minor component of the measured changes in NADH absorbance.

The free NADH concentration *[NADH]f* in the assay mix can be calculated using K_D_ for NADH binding to different subunits on GAPDH (Table 1):

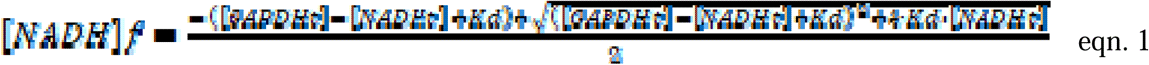

Where *[GAPDH]t* concentrations represent concentration in terms of NADH binding sites in GAPDH tetramers. [K_D_ is the GAPDH-NADH dissociation constant calculated from the fluorescence measurements (Table 1). The calculated free NADH concentration *[NADH]f* was used to calculate expected LDH activity in case of no channeling between LDH and GAPDH

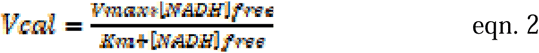

Vmax and K_M_ are the values for Michaelis-Menten constants for NADH oxidation reaction with LDH. All LDH assay mixtures were prepared with 630 μM pyruvate. Specific activities of phLDH, rmLDH, were 130 ± 15 and 430 ± 30 U/mg, respectively

### Sedimentation Velocity AUC Experiments

An-60-Ti rotor was used with three sample cells with quartz windows and charcoal-epon centerpieces with two sectors 12 mm optical path length. in each cell, 340 μL enzyme samples (6.0 µM) were loaded in one sector and 350 μL buffer in another. Two cells had individual enzymes, and the third cell had the enzyme mixture at the same loading concentration. Three sedimentation velocity experiments were measured in parallel by following absorbance at 280 nm. The first cell had a mixture of phLDH and byGAPDH, both enzymes in the concentration of 6 μM. The second cell had phLDH alone at concentrating of 6 μM, and the third cell had byGAPDH alone in the concentration of 6 μM. The highest protein concentration was determined by the maximal absorbance that can be measured by XL-A instrument (O.D. max=2.0). The experiments started with the prescans at 3000xg, and the sum of absorbances of the individual enzymes was compared to the absorbance of the mixture. The sedimentation profiles were measured for 7 hours at 40,000xg. Single scans with radial increments of 30 μm have been recorded every 240 sec. SEDNTERP was used with protein sequences to calculate molecular mass, partial specific volume, solvent densities, and viscosities ^38^. The measured sedimentation profiles were analyzed with SEDFIT program using nonlinear regression with *s*, and *Mr* as free fit parameters ^38^.

## Acknowledgments

DS & CHC work at the National Renewable Energy Laboratory operated by Alliance for Sustainable Energy, LLC, for the U.S. Department of Energy (DOE) under Contract No. DE-AC36-08GO28308. Funding was provided by the U.S. Department of Energy Office of Energy Efficiency and Renewable Energy (CHC) and Laboratory Directed Research and Development funds (DS). The views expressed in the article do not necessarily represent the views of the DOE or the U.S. Government. The U.S. Government retains and the publisher, by accepting the article for publication, acknowledges that the U.S. Government retains a nonexclusive, paid-up, irrevocable, worldwide license to publish or reproduce the published form of this work, or allow others to do so, for U.S. Government purposes Access to the high-performance computing was supported by the Ministry of Science and Education of the Republic of Croatia under the grant 533-19-15-0007 (Centre of Research Excellence for Data Science and Cooperative Systems). ZMS was a recipient of funds from Croatian Science Foundation’s project number O-1505-2015, funds from the University of Rijeka project number 511-12, and as a paid consultant for Jiva Pharmaceuticals MI, USA. The funders had no role in study design, data collection and analysis or decision to publish and preparation of the manuscript. ZMS and IO gratefully acknowledge services of NVIDIA CUDA Teaching Center and dr. Vedran Miletić. Gordan Janeš, Miroslav Puškarić, and dr, Draško Tomić provided help from the Center for Advanced Computing and Modeling. Ms. Eda Jardas has helped as a student researcher in the initial docking studies and *in silico* alanine scanning studies.

