## Supplemental for "Substrate Channeling *via* a Transient Protein-Protein Complex: The case of D-Glyceraldehyde-3-Phosphate Dehydrogenase and L-Lactate Dehydrogenase"

**Contents**

1.	Video: Interactions between rmLDH and rmGAPDH in presence and absence of NAD(H)	1

2.	Rigid-Body Docking of Protein-Protein Complex	3

3.	Replicates of all atom molecular dynamics simulation of binding interactions between different LDH and GAPDH molecules in absence and presence of NAD(H) molecules	8

4.	Coarse-grained molecular dynamics simulations of interaction between rmLDH (PDB:3H3F) and rmGAPDH (PDB:1J0X) in the absence of NAD(H)	11

5.	Numerical Simulations of NADH channeling in Enzyme Buffering Experiments in transient LDH-NADH-GAPDH complex	12

Supp. Fig. 11 (A-C). KinTek program was used for numerical simulation of LDH activity on its NADH substrate in the presence of a large excess of GAPDH [11].	15

Supp. Fig. 12 (A-C). Increase in *off-rates* for GAPDH-NADH complex can facilitate channeling in case of transient protein-protein interactions.	16

Supp. Fig. 13 (A-C). Changes in off-rates for GAPDH-NADH complex can reproduce experimentally observed differences between rmGAPDH and byGAPDH.	16

Supp. Fig. 14 (A-D). Simulation of Michaelis-Menten curves for LDH activity with its NADH substrate for “no channeling” and “channeling” paths in the presence of different concentrations of GAPDH	18

6. Full numerical description of enzyme buffering experiments	19

Supp. Table 1 A-B. NADH channeling from rmGAPDH-NADH (A) or byGAPDH-NADH (B) complex to rmLDH or phLDH (full numerical description of experiments presented in figure 7 and 8).	19

Supp. Table 2 A-B. NADH channeling from rmLDH-NADH (A) or phLDH-NADH (B) complex to rmGAPDH or byGAPDH	20

7.	Materials and Methods	21

8.	References	25

### 1. Video: Interactions between rmLDH and rmGAPDH in presence and absence of NAD(H)

**Supp. video 1.** All-atom molecular dynamics simulations of interactions between rmGAPDH (yellow ribbon, PDB:1J0X) and rmLDH (green ribbon, PDB:3H3F). The proteins are shown as ribbon models and as Van der Waals radii of each atom (transparent black shadows). The NAD(H) molecules are shown as black CPK models, and the surrounding water molecules and ions are not shown. The first frame in both simulations shows the two enzymes in the same position 5 Å apart facing each other with their NADH binding sites just as in Fig. 1A. The video on the left shows a gradual build-up of complex-forming interactions between rmGAPDH and rmLDH in the absence of NADH (Fig. 2A). The active site loop on LDH is visible in its open position as it is gradually latching to the GAPDH surface . The video on the right shows that GAPDH-NAD and LDH-NADH complexes can come in contact in different orientations but there is no gradual build-up of binding interactions. In LDH-NADH complex the active site loop is buried in its closed position . The simulation shows interaction build-up in 100 nanoseconds, with one frame recorded every nanosecond (Fig. 2). The simulation had 1.49 million atoms with 50 million steps and the step increase set at 2 femtoseconds. The box padding was set to 5 nm in each direction to provide large space for free diffusion.

**Supp. Video 2.** “*Morph Conformations*” utility was used to highlight the differences between NADH-free and NADH-bound structure for rmGAPDH (left, PDB:1J0X ) and for rmLDH (right, PDB:3H3F). In both presentations the reader is looking directly at the surfaces that form GAPDH-LDH complex. In GAPDH the nicotinamide portion of NAD(H) molecule binds in the cleft between the catalytic domain and NAD(H) binding domain and drives the compaction between the adjacent subunits . The compaction of the two GAPDH subunits will lead to the repositioning of its LDH binding sites by as much as 15.4 Å. In rmLDH, the transition from NADH-free to NADH-bound structures result in closing of the active site cleft. The closing is driven by repositioning of the active site loop (a.a. 96 to 105, yellow) and by twisting in the active site helix (a.a. 226 to 242, yellow). The active site loop is driven in its closed position by interactions between phosphate group on NADH and Arg 98 and by interactions between OH group on ribose and Arg 105 . The amino acids in the active site loop have the highest b-factor values (PDB:3H3F).

### 2. Rigid-Body Docking of Protein-Protein Complex

To gain a better understanding of how the GAPDH-LDH pose used in all-atom molecular dynamics relates to the general process of protein association, we undertook a docking study between the two proteins. Figure Supp 1 illustrates both proteins as they are posed in the docking “reference structure,” which we took from frame number 825 in the molecular dynamics simulation. We were not concerned per se with the process of protein-protein association, but rather how representative the structure used in atomistic molecular dynamics is of the initial binding ensemble one might expect. Figure Supp 2 illustrates the 100 lowest energy structures found considering only electrostatic and van der Waals forces between the two partners. To create an interpretable visualization, LDH was encoded as arrow pairs, as explained in Methods. Notably, with this basic energy expression, one observes a fairly uniform distribution of LDH docking around the GAPDH model. There are two objections to be made. First, that GAPDH is a tetramer, and as posed in the figures the dimeric model “back side” (facing toward the right of the figures) would be occupied by partner GAPDH subunits. One can safely dismiss LDH poses around the back side of GAPDH as an artifact of approximate modeling. Second, the energy model takes no account of desolvation contributions to binding. To address this point, figures Supp 3 and Supp 4 show the 100 lowest energy structures when incorporating two alternative desolvation models, Atom Contact Energy or Optimal Docking Area . Two pleasing developments are seen relative to the solvation-free models. First, there is a clear depopulation of poses around the “top” and “bottom” of GAPDH, which would be solvent-exposed but unproductive in catalysis. Second, there is a pronounced clustering of poses that are obviously artifactual given the dimeric model. Thus, while LDH favorably binds to the internal surfaces of the GAPDH tetramer in the model world, these poses are irrelevant to the physical world by virtue of occlusion from the unmodeled GAPDH subunits.

Nevertheless, what figures Supp 3 and Supp 4 show is that the reference structure is by no means representative of what would be expected as an initial encounter complex (which is expected, given that the “open” conformation of LDH involved in actual binding was not modeled). Furthermore, rigid body docking will not account for sidechain or backbone motions that can develop full binding energy. Finally, the observation suggests that the association of GAPDH and LDH into the proposed channeling-active reference structure is not a “just-so” story—the binding conformation explored does not represent a dramatic, deep energetic well into which the partner enzymes fall. Rather, this evidence suggests that evolution of a channeling-active structure is likely dynamic, and that channeling need not be tightly coupled to a particular structure, but could involve a more “leaky” model where transfer of NADH from GAPDH to LDH may benefit both from specific interactions, as well as random diffusion around approximately productive complexes.

To further document the relationship of the reference complex to the docking ensemble, the docked structures were filtered according to distance constraints that would recover the reference conformation. Figure Supp 5 illustrates the 11 structures that satisfied these constraints, and Figure Supp 6 shows the relationships among the three different energetic models explored. The energetic relationships were then analyzed for the resulting structures, for the three energy models considered. ES+vdW (the base energetic model) has very little relationship to the desolvated models, as seen from the extensive intersections among the blue lines. However, both desolvation models capture a more similar ordering and energetic range. What falls out of this analysis is that the reference structure is neither too high nor too low in energy (within the limitation of a rigid-body approximation), and that desolvation of the protein interaction surface is likely an important component of complex formation, given that the second worst (highest energy) of the ES+vdW complexes becomes the best under both desolvation models.

**Table Supp 1. Atom type mapping among NADH AMBER, ACE, and ODA types.**

| PDB Atom Name | AMBER Type | ACE Type | ODA Type |
| --- | --- | --- | --- |
| O1N | O2 | DOD | O- |
| O2N | O2 | DOD | O |
| O1A | O2 | DOD | O- |
| O2A | O2 | DOD | O |
| O3 | OS | DOD | OH |
| PN | P | DOD | S |
| PA | P | DOD | S |
| C6N | CA | FCZ | Carom |
| C5N | CA | FCZ | Carom |
| C3N | CA | FCZ | Carom |
| C2N | CA | FCZ | Carom |
| C6A | CA | FCZ | Carom |
| C5A | CB | FCZ | Carom |
| C4A | CB | FCZ | Carom |
| C8A | CK | FCZ | Carom |
| C2A | CQ | FCZ | Carom |
| C4N | CT | FCZ | C |
| C1D | CT | FCZ | C |
| C2D | CT | FCZ | C |
| C3D | CT | FCZ | C |
| C4D | CT | FCZ | C |
| C5D | CT | FCZ | C |
| C5B | CT | FCZ | C |
| C4B | CT | FCZ | C |
| C3B | CT | FCZ | C |
| C2B | CT | FCZ | C |
| C1B | CT | FCZ | C |
| N1N | N* | HNE | N |
| N9A | N* | HNE | N |
| N7A | NB | HNE | N |
| N1A | NC | HNE | N |
| N3A | NC | HNE | N |
| N6A | N2 | KNZ | N |
| C7N | C | NND | C |
| N7N | N | NND | N |
| O7N | O | NND | O |
| O2D | OH | SOG | OH |
| O3D | OH | SOG | OH |
| O3B | OH | SOG | OH |
| O2B | OH | SOG | OH |
| O4D | OS | SOG | OH |
| O5D | OS | SOG | OH |
| O5B | OS | SOG | OH |
| O4B | OS | SOG | OH |

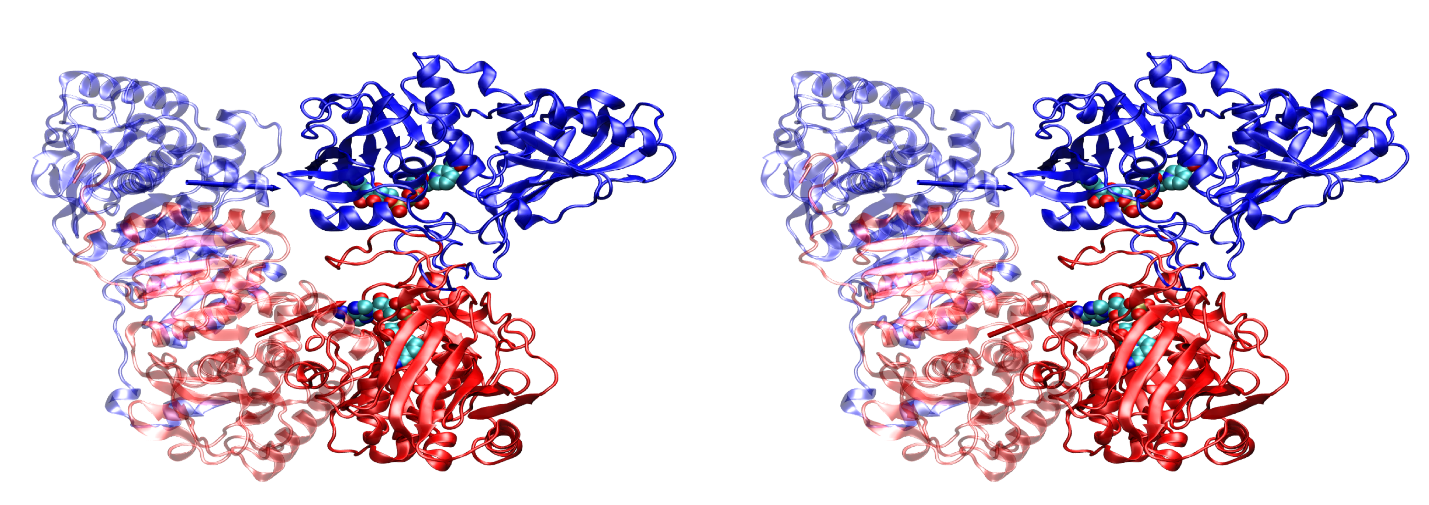

**Supp. Fig. 1. Walleyed stereo view of rmLDH (PDB:3H3F) and rmGAPDH (PDB:1J0X) in the absence of NAD(H).** The interacting subunits in rmLDH-rmGAPDH complex are shown in the same orientation as in figure 1. Arrows visible in the rmLDH structure extend from Arg 168 atom NH1 to Lys 242 atom NZ, and represent the approximate orientation of the substrate binding site to the surface. These arrows are used in figures below to denote LDH orientation relative to central rmGAPDH. Blue denotes LDH chain D or GAPDH chain Q; red, LDH chain C or GAPDH chain P. The figures show rmLDH-rmGAPDH complex that forms in the absence of NAD(H), and for orientation purposes the NAD(H) binding sites are marked with molecules represented with van der Waals spheres.

| 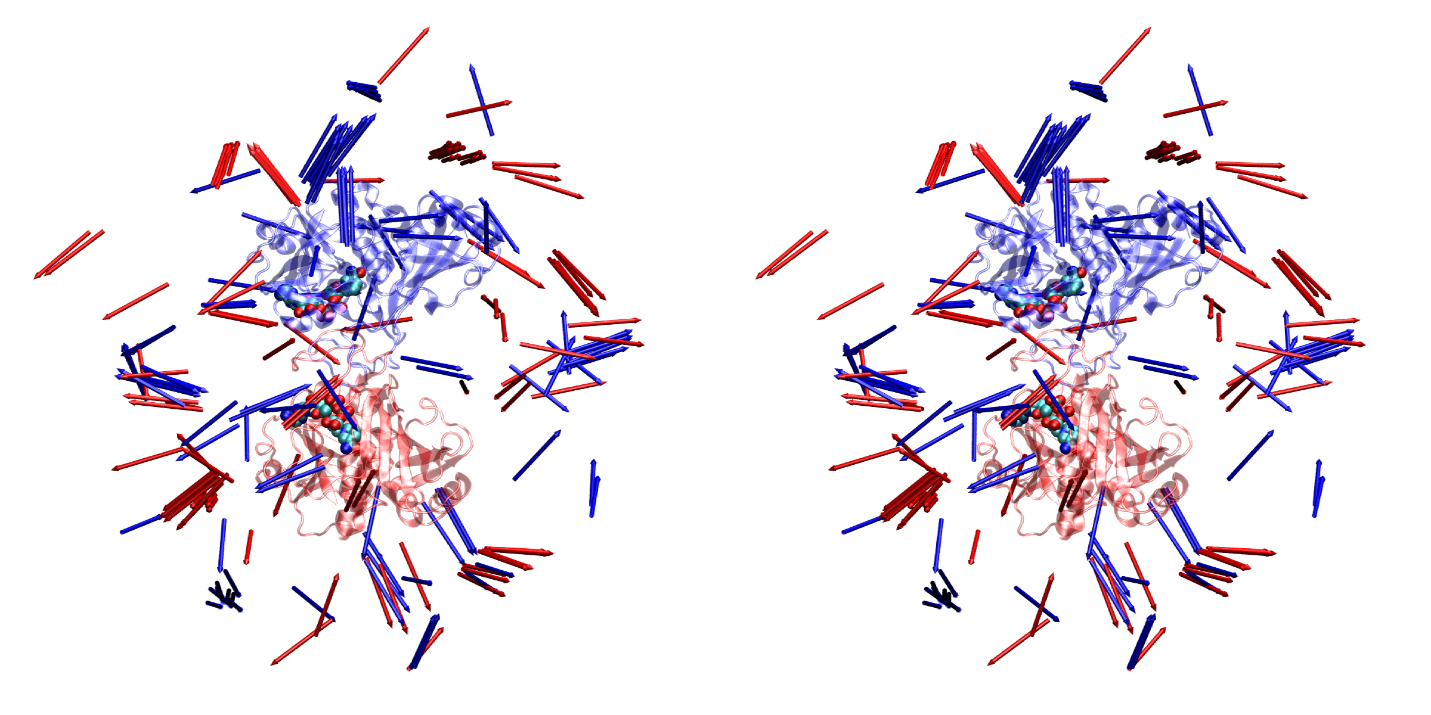 | **Supp. Fig. 2.** Walleyed stereo view of top 100 lowest energy docking configurations of LDH around GAPDH. Energy includes only electrostatic and van der Waals forces. Color coding, GAPDH orientation, NADH representation, and arrow definitions are as in Supp. Fig. 1. |
| --- | --- |

| 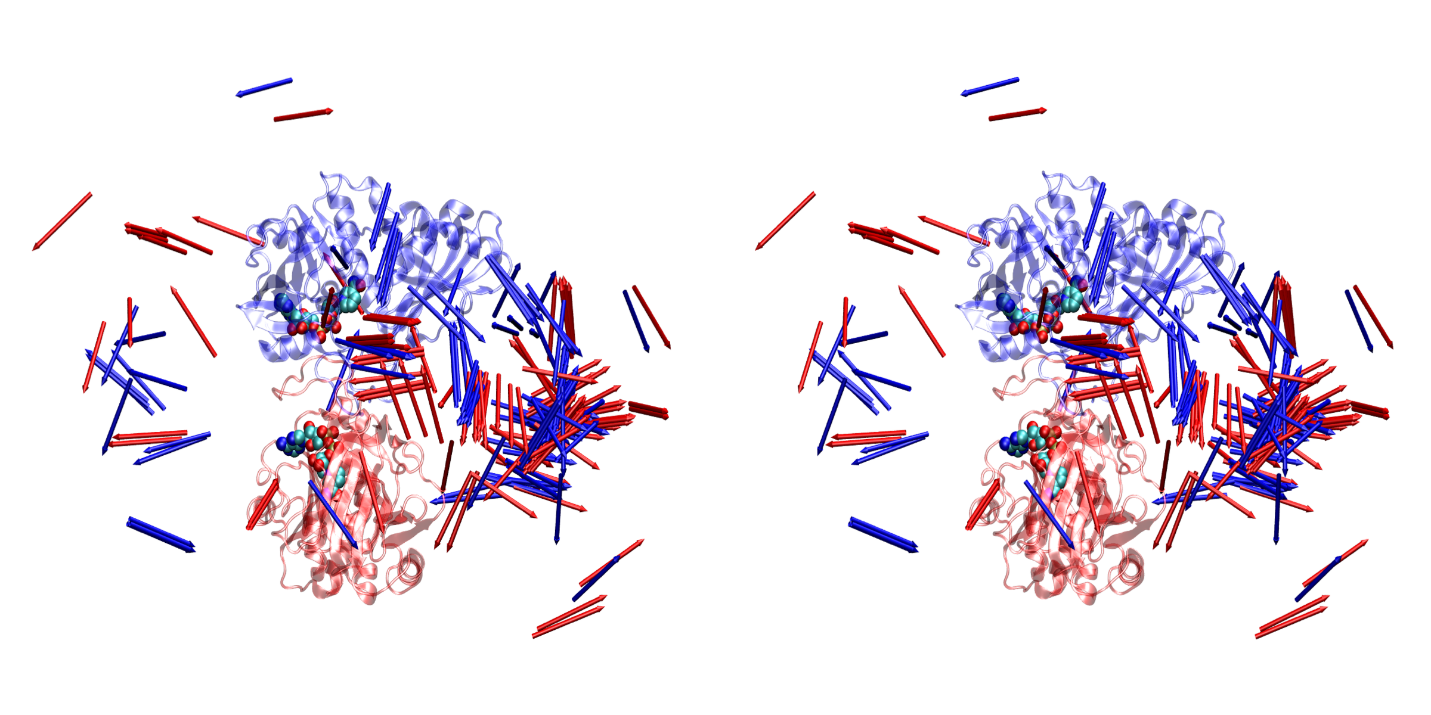 | **Supp. Fig. 3.** Walleyed stereo view of top 100 lowest energy docking configurations of LDH around GAPDH. Energy includes electrostatic and van der Waals forces, plus that from the ACE desolvation model . Color coding, GAPDH orientation, NADH representation, and arrow definitions are as in Supp. Fig. 1. |
| --- | --- |

| 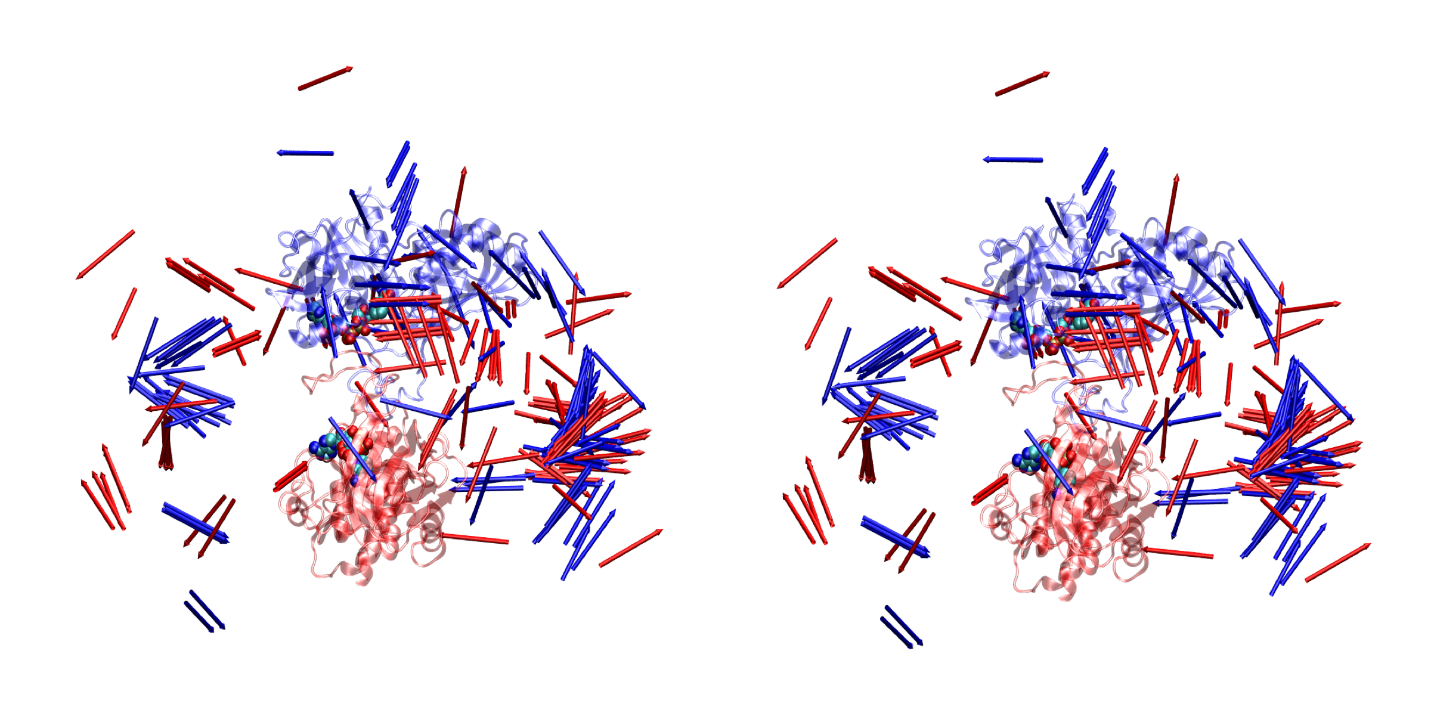 | **Supp. Fig. 4.** Walleyed stereo view of top 100 lowest energy docking configurations of LDH around GAPDH. Energy includes electrostatic and van der Waals forces, plus that from the ODA desolvation model . Color coding, GAPDH orientation, NADH representation, and arrow definitions are as in Figure Supp 1. |
| --- | --- |

| 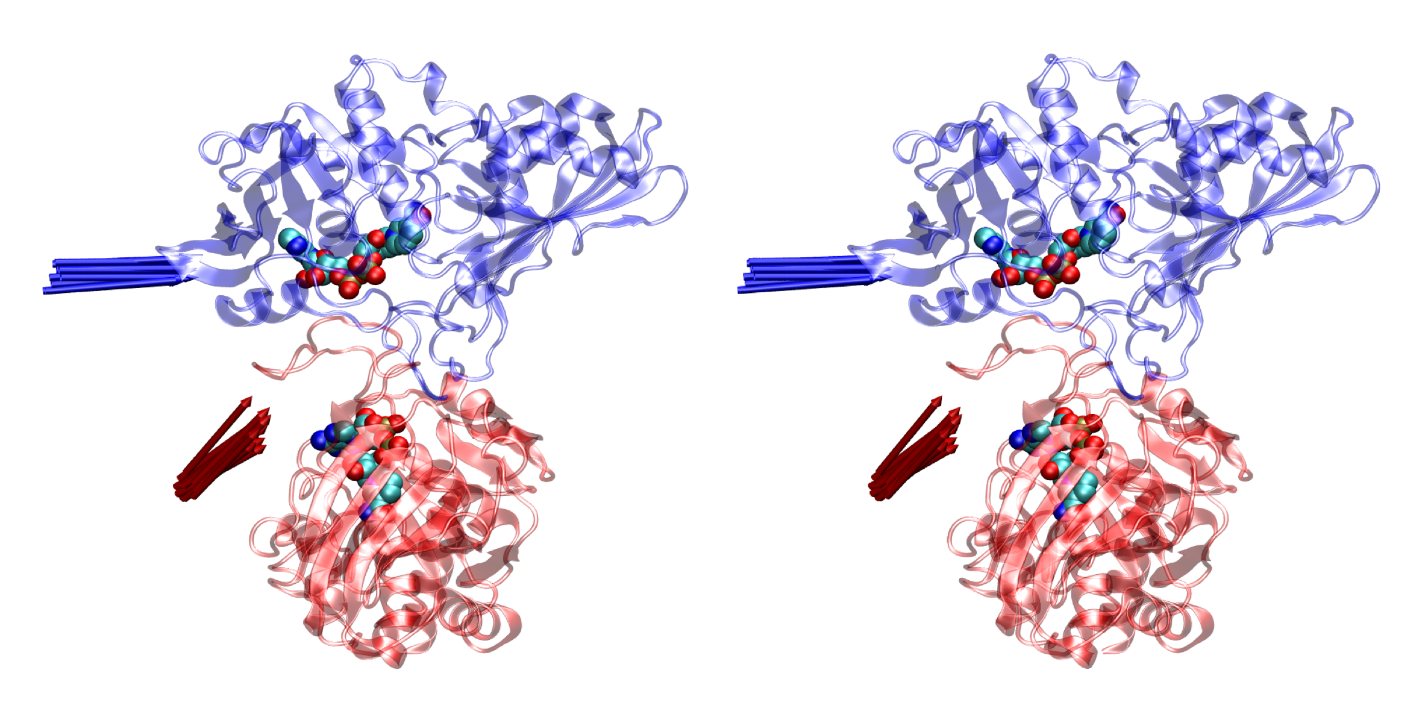 | **Supp. Fig. 5.** Walleyed stereo view of top 11 docking configurations of LDH around GAPDH satisfying distance constraints between LDH chain D and GAPDH chain Q. Color coding, GAPDH orientation, NADH representation, and arrow definitions are as in Supp. Fig. 1. |
| --- | --- |

| 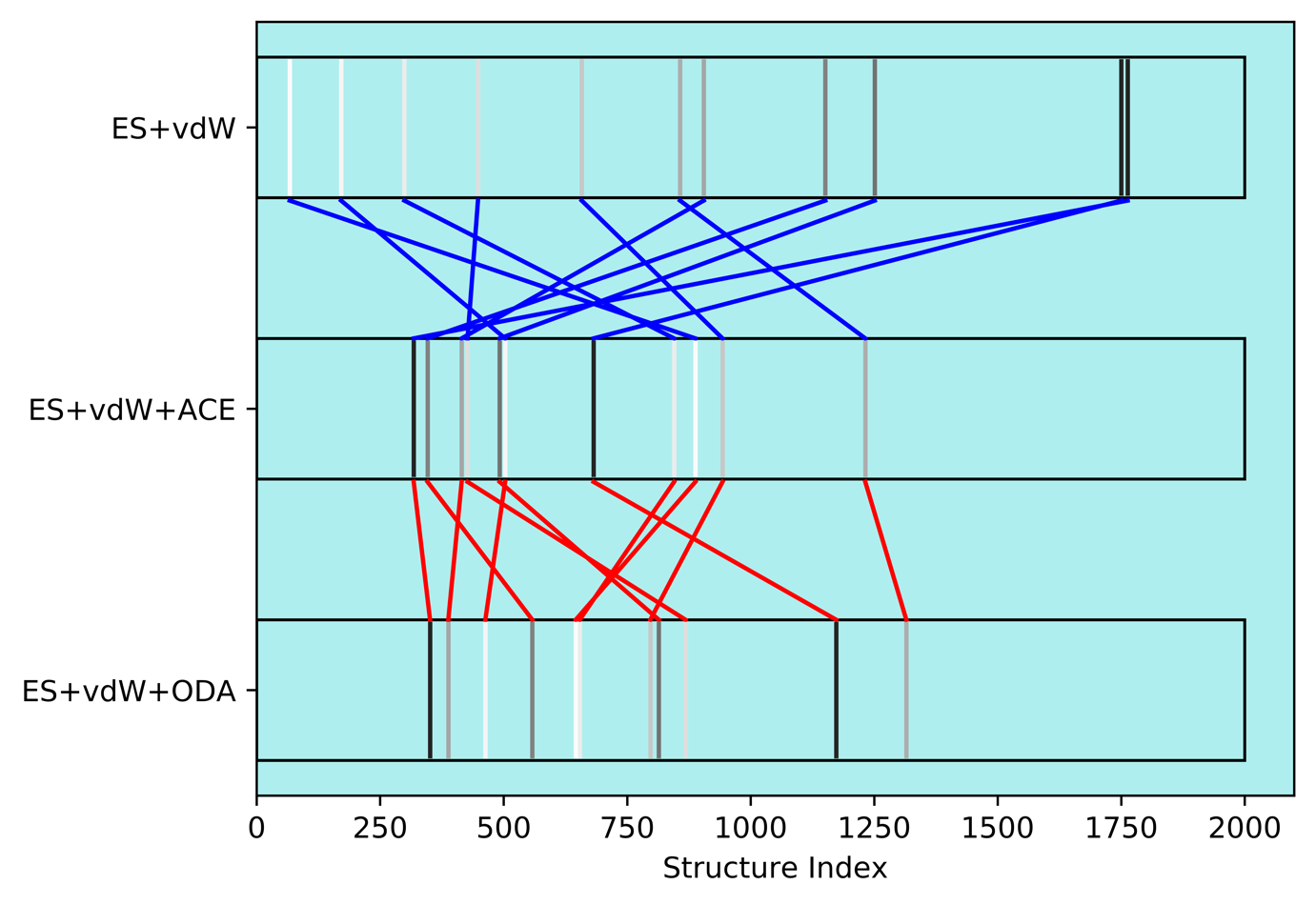 | **Supp. Fig. 6.** Correlation plot showing relationships among the 11 distance-filtered docked complexes, under the three different energy models explored. Throughout, each complex is represented as a line segment, gray-scaled according to its electrostatic + van der Waals (ES+vdW) energy. Positions along the Structure Index are proportional to energies under the corresponding model. Relationships between ES+vdW energy and that augmented by the ACE desolvation model (ES+vdW+ACE) are highlighted with blue lines; between ES+vdW+ACE and the alternative ODA desolvation energy (ES+vdW+ODA) with red lines. |
| --- | --- |

### 3. Replicates of all atom molecular dynamics simulation of binding interactions between different LDH and GAPDH molecules in absence and presence of NAD(H) molecules

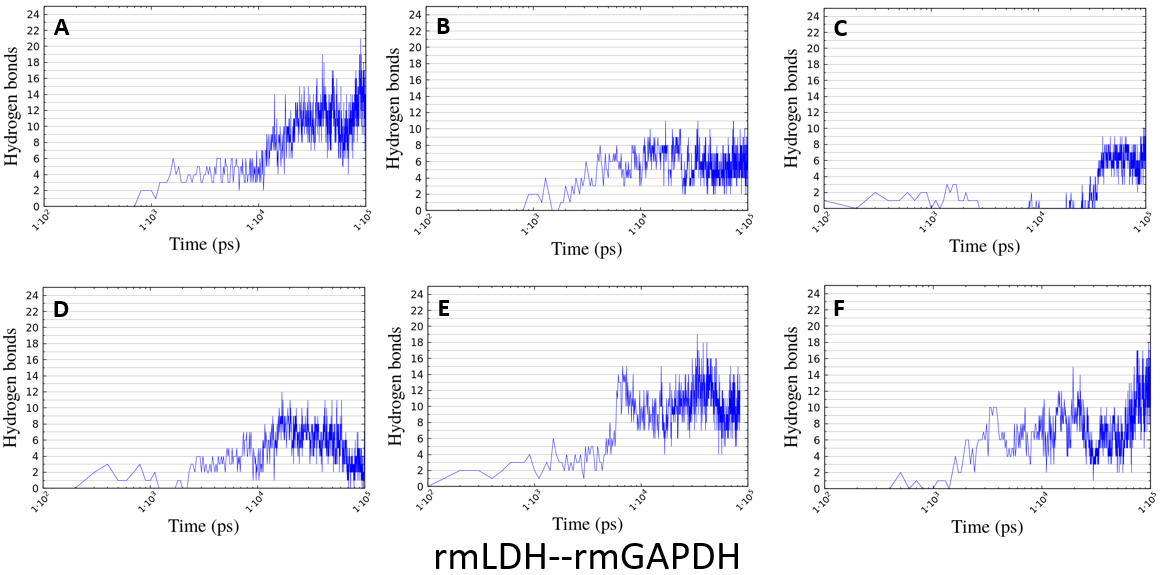

**Supp. Fig. 7 (A-F). Replicates of all atom molecular dynamics simulation of binding interactions between rmLDH (PDB:3H3F) and rmGAPDH (PDB:1J0X) in the absence of NAD(H).** The figure shows how small changes in the initial docking orientation and the position of the flexible parts on the protein surface can affect the build-up of binding interactions. The gradual build-up in binding interactions is a result of wobbling between the two protein complexes and mobility of the loops on the protein surfaces. Panels A, E, F, show gradual build-up of binding interactions as the energy of interaction goes through a series of local energy minima until the most stable complex is achieved. Panels B, C, D, show that in some simulations the energy of interaction can stay entrapped in local energy minima, and the gradual build-up of binding interactions never happens.

The gradual buildup of binding interactions was calculated using built-in functions in program GROMACS (cut-off values set at 3.5 A and 25 degrees) . All simulations started with the two proteins 5 Å apart facing each other with their NAD(H) binding sites . Thus, there are no binding interactions until the two proteins collide driven by random diffusion. Simulations are presented in a logarithmic scale to show the gradual build-up in binding interactions throughout the 50 million simulation steps (100 nsec).

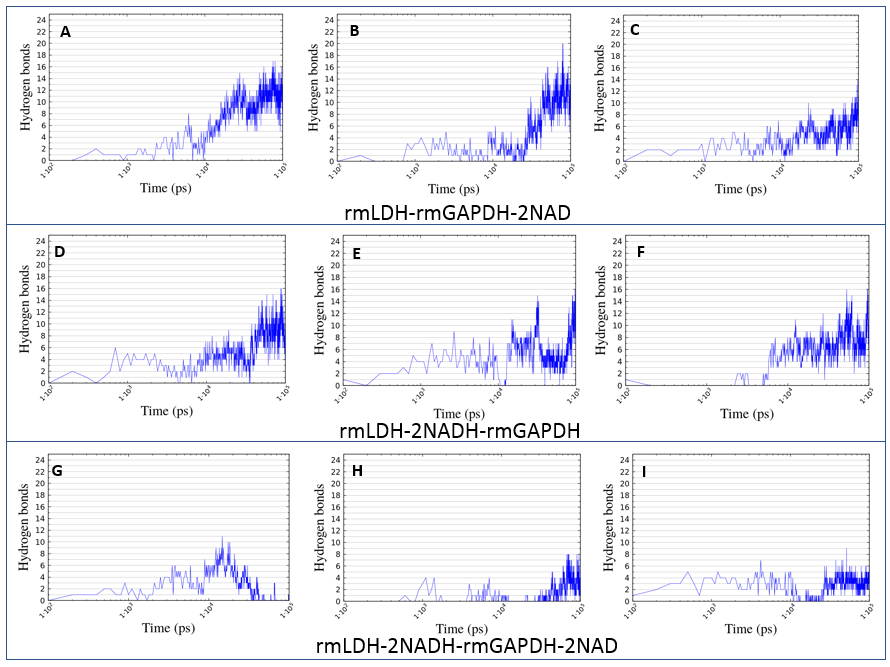

**Supp. Fig. 8 (A-I). Replicates of all-atom molecular dynamics simulation of binding interactions between rmLDH (PDB:3H3F) and rmGAPDH (PDB:1J0X) with different NAD(H) complexes.**

The gradual buildup of binding interactions was calculated using built-in functions in program GROMACS (cut-off values set at 3.5 A and 25 degrees) . All simulations started with the two proteins 5 Å apart facing each other with their NAD(H) binding sites . Thus, there are no binding interactions until the two proteins collide driven by random diffusion. Simulations are presented in a logarithmic scale to show the gradual build-up in binding interactions throughout the 50 million simulation steps (100 nsec). The presentation shows that saturation with NAD(H) does not support a gradual build-up of rmLDH-rmGAPDH complex (G-I), the two proteins fall apart (G) or form only few random collisions (H-I).

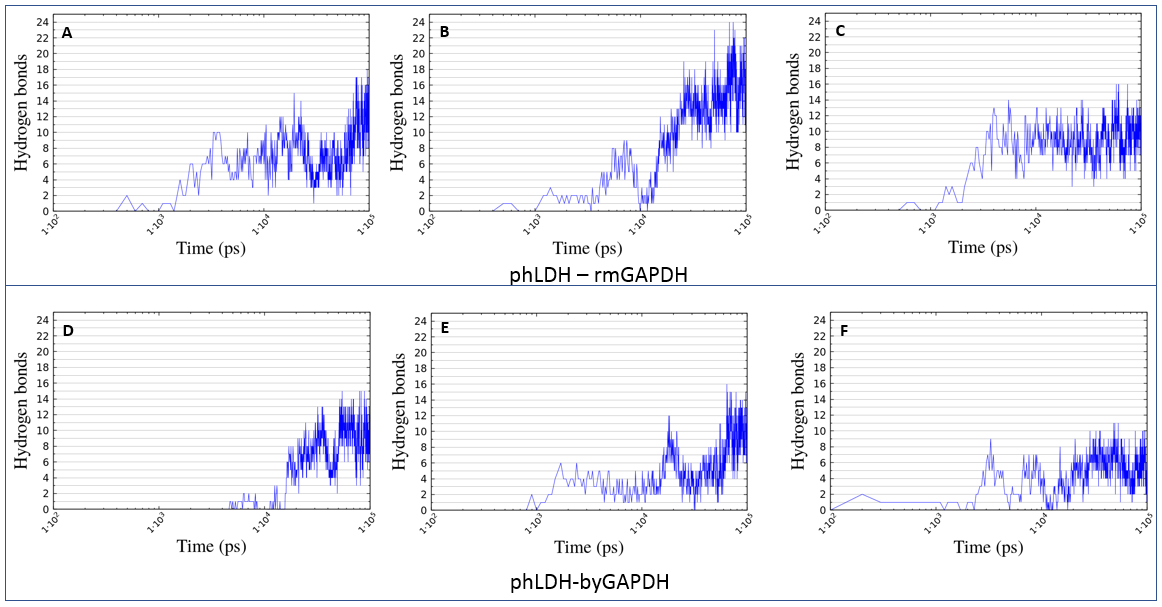

**Supp. Fig. 9. Replicates of all atom molecular dynamics simulation of binding interactions:**

**(A-C) between porcine heart LDH (PDB:5LDH) and rabbit muscle GAPDH (PDB:1J0X)**

**(D-F) between rabbit muscle LDH (PDB:5LDH) and baker’s yeast GAPDH (PDB:3PYM).**

The gradual buildup of binding interactions was calculated using built-in functions in program GROMACS (cut-off values set at 3.5 A and 25 degrees) . All simulations started with the two proteins 5 Å apart facing each other with their NAD(H) binding sites . Thus, there are no binding interactions until the two proteins collide driven by random diffusion. Simulations are presented in a logarithmic scale to show the gradual build-up in binding interactions throughout the 50 million simulation steps (100 nsec). The presentation shows that there is a subtle but statistically significant difference in the number of binding interactions in phLDH-rmGAPDH versus rmLDH-byGAPDH complex. rmGAPDH (PDB: 1J0X) and byGAPDH (PDB: 3PYM, isozyme 1) have 65.5% sequence identity and 84.8% similarity, the corresponding structures can overlap with RMSD value of 0.556 Å. phLDH has 75% sequence identity and 93.1% sequence similarity with rmLDH. The corresponding structures can overlap with RMSD value of 1.73 Å (5LDH and 3H3F ),

### 4. Coarse-grained molecular dynamics simulations of interaction between rmLDH (PDB:3H3F) and rmGAPDH (PDB:1J0X) in the absence of NAD(H)

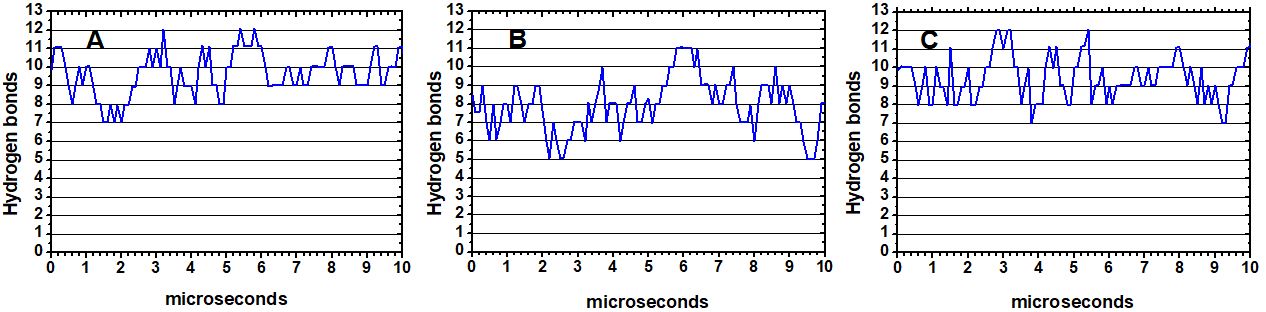

**Supp. Fig. 10 (A-C). Replicates of coarse-grained molecular dynamics simulation of binding interactions between rmLDH (PDB:3H3F) and rmGAPDH (PDB:1J0X).** The simulations started with the two proteins 5 Å apart facing each other with their NAD(H) binding sites just as in all-atom simulations (Supp. Fig 7) . The number of binding interactions was calculated using built-in functions in program GROMACS (cut-off values set at 3.5 A and 25 degrees) . The binding interactions form within the first 0.1 µsec of simulation, and complex remains stable throughout 10 µsec. The coarse-grained simulation can activate the motions between the NAD(H) binding domain and the catalytic domain in both LDH and GAPDH that cannot be activated in the all-atom simulations (Supp Fig 7). Those motions are a part of the apo-enzyme holo-enzyme transition and have a direct effect on the stability of the complex (Supp. Video 1-2). Thus the motions that drive apo-enzyme holo-enzyme transitions correlate with the oscillation in the number of binding interactions observed in the coarse grained simulations.

### 5. Numerical Simulations of NADH channeling in Enzyme Buffering Experiments in transient LDH-NADH-GAPDH complex

The enzyme assays with two interacting enzymes that share a common substrate can be challenging to design and interpret . We used numerical simulations to prepare for the assay design and the data interpretation (Supp. Fig. 11-14). The simulations can also help us to avoid some of the misunderstandings that have plagued many of the earlier studies of substrate channeling .

Using numerical simulations, we found that model mechanism that depicts NADH channeling from GAPDH to LDH *via* transient protein-protein complex can reproduce the key experimental features observed in assays with different LDH-GAPDH pairs (Fig 4-7 vs. Supp. Figs. 11-14). Simulations show that substrate channeling is a regulated molecular mechanism in which partner enzymes can affect each other activity while sequestering the shared substrate. Most of the presented conclusions are likely to be valid for all cases of substrate channeling *via* transient protein-protein interactions .

*Enzyme buffering experiments and substrate channeling in transient protein-protein interactions*. We analyzed LDH activity with NADH substrate in the presence of a large excess of GAPDH (Figs. 7-9). Such measurements can mimic LDH activity in the cytosol, where the majority of NAD(H) molecules are bound to GAPDH, the most abundant NADH dehydrogenase in cells . Following the presented LDH-GAPDH complex (Fig. 1), we have devised a molecular mechanism that consists of 9 different interactions with 12 different sets of rate constants (Supp. Fig. 11A). Different experimental conditions can be analyzed using different sets of on and off rate constants and various concentrations of LDH, GAPDH and NADH (Supp. Fig. 11A). The rate constants used in our simulations are within an order of magnitude equal to the measured rate constants, which is consistent with experimental variability in such measurements .

LDH activity in the case of *no-channeling* between GAPDH and LDH can be simulated by setting to zero rate constants for steps 5 to 12 in the channeling reaction diagram (Supp. Fig. 11A). The simulations show that in the presence of a large excess of GAPDH, the majority of NADH molecules are bound to GAPDH-NADH complex (Supp. Fig. 11A). Thus, the majority of LDH molecules cannot bind NADH and remain inactive even at the saturating NADH concentrations (NADH=40 µM). Repeated simulations with different LDH, GAPDH, and NADH concentrations showed that the majority of LDH molecules will be in NADH-free-form in all cases where GAPDH concentration is high enough to drive the free NADH concentration well below its *Km* constant for LDH (Supp. Fig. 11 A-B). In such conditions, LDH activity is controlled by diffusion limited formation of LDH-NADH complex.

Panel C in Supp. Fig. 11. shows the same simulation as panel B, except that this simulation included different interactions between LDH and GAPDH that can take place in the case of channeling. We know very little about the channeling interactions. Thus, for steps 5 to 12, we decided to analyze the most challenging experimental situation, the weak transient protein-protein interactions that cannot be readily observed in routine studies of protein-protein interactions . In the case of transient interactions, the LDH-GAPDH complex can form relatively fast, but it is also relatively short-lived .

A significant simplification in the selection of the rate constants for channeling reaction comes from the rules of thermodynamic cycles . The rules state that changes in free energy of interaction do not depend on the pathway taken, but only on the initial and the final state. In our case that means *ΔG1+ ΔG10= ΔG5+ ΔG8* or *Kd1•Kd10 = Kd5•Kd8* . For example, if there is a change in NADH binding affinity in LDH-GAPDH complex relatively to free LDH (*Kd1 vs. Kd8*), then there should be a compensatory change in LDH binding affinity for GAPDH relative to GAPDH binding affinity of LDH-NADH complex (*Kd5 vs. Kd10*). The same logic applies for the second cycle, NADH binding to LDH-GAPDH complex (*Kd7 and Kd8*) and NADH transfer rates from LDH-(GAPDH-NADH) to (LDH-NADH)-GAPDH (*Kd9*). The third cycle requites that *Kd3•Kd5•Kd12 = Kd2•Kd10•Kd11* (Supp. Fig. 11A).

We show an example of simulation where observed LDH activity with the channeling path included is 2.7-fold higher than the corresponding reaction with no channeling (Supp. Fig. 11C vs. 11B). The simulation shows transient interactions where *Kd1•Kd10 = Kd5•Kd8* = 1000 μM^2^. Dissociation constants for LDH-GAPDH, LDH-(GAPDH-NADH) were set to 200 μM and (LDH-NADH)-GAPDH were set to 100 μM (Supp. Fig. 11A). These constants represent transient protein-protein interactions that are too weak to be detected by routine protein-protein interaction studies. Following the rules of thermodynamic cycles selected choices for *Kd5* and *Kd10* to imply that the dissociation constant for (LDH-NADH) complex *Kd1* has to be two times bigger than the NADH dissociation constant for GAPDH-LDH complex *Kd8*. Such an assumption can be justified by the bigger positive fields around NADH binding sites in LDH-GAPDH complex (Fig. 1). The NADH transfer rate within (GAPDH-NADH)-LDH complex was set to be equal to the off-rates for the GAPDH-NADH complex (Supp. Fig. 11A, *k9* = *k-4*). In other words, in the presented mechanism, LDH binding to GAPDH-NADH complex does not affect the binding interactions between GAPDH and NADH.

Simulations showed that the channeling and the free-diffusion paths always take place in parallel. The two paths cannot be separated experimentally. However, the simulations can be used to show that the rates in the “no-channeling” path are equal to the *calculated free diffusion rates* (methods eqns. 2-3). The *calculated free diffusion rates* are easy to calculate in experiments using the constants that can be measured with high accuracy (Fig. 6). Accordingly, the substrate channeling can be detected in enzyme buffering experiments if the *observed LDH activity* is higher than the *calculated free diffusion LDH activity* (methods eqns. 2-3).

*Substrate channeling and dissociation rates (koff) for GAPDH-NADH complex Supp. Fig. 12)*. We want to understand to what extent GAPDH binding affinity for NADH can affect the channeling path (*Kd* for steps 4,7 and 9 in Supp. Fig. 11). More specifically we analyzed *off*-rates for GAPDH-NADH complex (steps *k-4*, *k-7*, and *k9* in Supp. Fig. 11). The factors that control off-rates for GAPDH-NADH complex are particularly interesting because they are also likely to affect the channeling rates. i.e., NADH transfer from (GAPDH-NADH)-LDH to GAPDH-(LDH-NADH) complex (Supp. Fig 11A, rate *k-4* vs. rate *k9*).

Simulations presented in Supp. Fig. 11 have been repeated with dissociation constants for GAPDH-NADH complex set to 5, 10, 20, 30, 40 sec-1 (steps: *k-4*, *k-7*, and *k9*). Simulations showed that increase in *off*-rates for GAPDH-NADH complex leads to decrease in concentration of (GAPDH-NADH)-LDH and GAPDH-NADH complex and increase in concertation of GAPDH-(LDH-NADH), GAPDH-LDH and LDH-NADH complex and in free NADH concertation. (Supp. Fig. 12 A-B). The result is a hyperbolic increase in the ratio between the *observed LDH activity* and the *calculated-free-diffusion rate* activity (Supp. Fig. 12 C).

The presented results have several significant conclusions:

1. In the case of transient protein-protein interactions the channeling depends on an overlap in timing of two events: formation of LDH-(NADH-GAPDH) complex and NADH dissociation from GAPDH within (GAPDH-(NADH)-LDH) complex. This conclusion is likely to be valid for all cases of substrate channeling via transient protein-protein interactions .

2. The off-rates for the protein-substrate complex are directly proportional to the complex dissociation constant Kd (Kd=koff/kon). Thus, to first approximation differences in the substrate binding affinity Kd can be used to predict chances of channeling for different protein pairs. That indicates that channeling can be affected by the negative cooperativity which regulates NAD(H) binding affinity for different subunits on GAPDH , or there could be difference in channeling between NADH and NAD substrates , or there could be differences in channeling from GAPDH to LDH relative to the channeling from LDH to GAPDH .

*Increase in the off-rates for GAPDH-NADH complex can explain experimentally observed differences between rmGAPDH and byGAPDH* (*Figs. 7 and 8 vs. Supp. Fig. 13*). LDH activity was analyzed in the presence of increasing apoGAPDH concentrations (Supp. Fig. 13). The simulations can easily show that increase in apoGAPDH concertation at fixed NADH concertation favors channeling because it leads to increase in concertation of GAPDH-NADH, (GAPDH-NADH)-LDH and GAPDH-LDH complexes and decrease in concertation of free NADH, free LDH, and LDH-NADH complex.

To analyze the differences in NADH binding affinity between byGAPDH and rmGAPDH we have repeated the simulations with increasing GAPDH concentrations with the *off*-rates for GAPDH-NADH complex set to 40 sec-1 and 5 sec-1 (*k-4, k-7*, and *k9* in Supp. Fig. 11 A). The simulations showed that increase in the *off*-rates will increase both the *observed LDH activity* and the *calculated free-diffusion activity* (Supp. Fig. 13 A-B). The higher ratios between the *observed LDH activity* and the *calculated free-diffusion activity* (Supp. Fig. 13 A-B) show that the higher *off*-rates have bigger effect on the channeling path than on the free-diffusion path.

The presented simulations show that the model mechanism with higher *off*-rate (higher *Kd* constants) for GAPDH-NADH complex is consistent with the experimentally observed differences between byGAPDH and rmGAPDH (Fig. 7 and 8).

*Modulation of channeling reaction path by decrease in total NADH concertation (Supp. Fig 14)*. Simulations can show that a decrease in total NADH concertation in the assay mix leads to a decrease in relative concertation of LDH-NADH complex and increase in concentration of different LDH-GAPDH complexes. The result is increase in contribution from the channeling reaction path and a decrease in contribution from the free-diffusion path (Supp. Fig. 14 A-B). Accordingly, the highest ratio between the *observed LDH activity* and the *calculated free-diffusion activity* can be seen when the lowest total NADH concertation is combined with the highest GAPDH concentration (Supp. Fig. 14 C). This observation is consistent with experiments which showed that that highest substrate channeling can be observed at lowest NADH concentrations (Fig 7-8 and ).

One of the main questions in metabolism studies is how channeling can affect the Michaelis-Menten constants. We found that GAPDH acts as a competitive inhibitor of LDH activity (Supp. Fig. 14 B and D). Simulations can show that inhibition is due to different LDH-GAPDH complexes that can slow down the catalytic cycle by adding extra steps to the channeling path relative to the free diffusion path (Supp. Fig. 11A). Studies of cellular metabolism usually describe channeling as a process that leads to more efficient metabolism due to sequestration of the shared metabolites. Our results indicate that transient protein-protein interactions can regulate channeling and cellular metabolism by a molecular mechanism that is more versatile than a simple sequestration of the channeled substrate (Supp. Figs. 11-14).

| 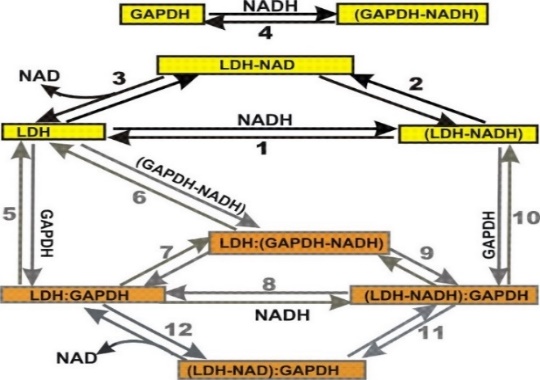  **A** | *on* constants | *off* constants | Dissociation  Constants |
| --- | --- | --- | --- |
|  | 10^6^ M^-1^ sec^-1^ | sec^-1^ | μM |
|  | k1= 2 | k-1= 20 | Kd1= 10 |
|  | k3= 0.1 | k-3= 40 | Kd3= 400 |
|  | k4= 5 | k-4= 40 | Kd4= 8 |
|  | k5= 20 | k-5= 4000 | Kd5= 200 |
|  | k6= 50 | k-6= 10000 | Kd6= 100 |
|  | k7= 2 | k-7= 40 | Kd7= 20 |
|  | k8= 4 | k-8= 20 | Kd8= 5 |
|  | k10= 2.5 | k-10= 250 | Kd10=100 |
|  | k12=0.1 | k-12=40 | Kd12=400 |
|  | rate, sec-1 | rate, sec-1 | Process |
|  | k2= 1000 | k-2= 5 | Turnover rate |
|  | k9= 40 | k-9= 20 | Channeling |
|  | k11=1000 | k-11=5 | Turnover rate |
|  | LDH Km NADH= 18 μM (calculated from simulation)  LDH Vmax = 34 sec-1 (calculated from simulation) | | |
|  | Component concentrations: | | |
|  | [LDH]= | 1 micro M | |
|  | [GAPDH]= | 400 micro M | |
|  | [NADH]= | 40 micro M | |
| 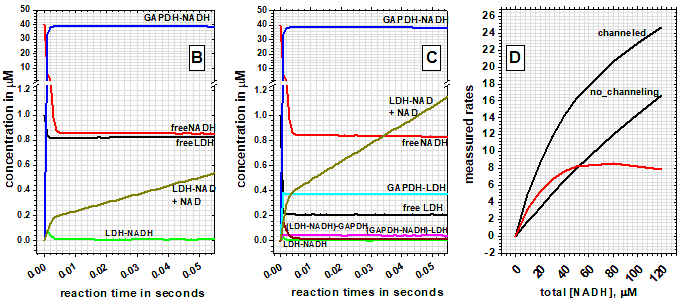 | | | |

#### **Supp. Fig. 11 (A-C). KinTek program was used for numerical simulation of LDH activity on its NADH substrate in the presence of a large excess of GAPDH .**

**(A)** Substrate channeling from GAPDH-NADH complex to LDH is illustrated schematically. Yellow boxes show steps in LDH reaction with free NADH, and orange boxes show steps in LDH reaction with GAPDH-NADH complex. Different reaction steps are labeled with numbers 1 to 12. Different experimental conditions can be analyzed using different sets of on and off rate constants and different LDH, GAPDH and NADH concentrations The values for different constant were chosen based on the available experimental data and by following the rules of thermodynamic cycles .

(**B**) The panel shows the LDH reaction with free NADH, without any interaction between LDH and GAPDH or GAPDH-NADH complexes (i.e., no channeling). The rate constants for steps 1 to 4 are listed in the table, while the rate constants for “the channeling steps” 5 to 12 were set to zero. The simulation shows that in these conditions about 97% of NADH is bound to GAPDH, and about 96% of LDH is in free form. The reaction time is adjusted so that the pre-steady state burst and the steady-state activity are visible . The simulations can be used to show that “no-channeling” activity is equal to the calculated free diffusion activity (methods eqns. 2-3). The related *Km* and *Vmax* constants can be calculated by repeating the simulation at different NADH concertation.

(**C**) The panel shows the same situation as panel B, except that this time the simulation included different interactions between LDH and GAPDH or LDH and GAPDH-NADH complex (steps 5 to 12). The observed LDH activity is 2.7 times higher than in the panel B. Event at GAPDH concertation at 400 µM about 20% of LDH molecules are free. Only about 1% of GAPDH molecules are present as different LDH-GAPDH complexes. The simulation shows that in enzyme buffering experiments significant NADH channeling can happen even when LDH-GAPDH interaction can be difficult to detect.

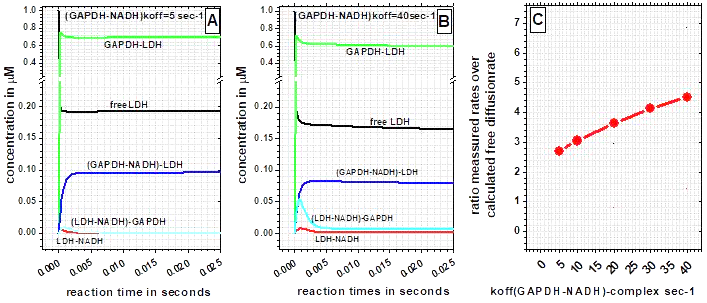

#### **Supp. Fig. 12 (A-C). Increase in *off-rates* for GAPDH-NADH complex can facilitate channeling in case of transient protein-protein interactions.**

**(A-B)** All rate constants and concentration in these simulations were the same as in Supp. Fig 11 except that the dissociation rates for GAPDH-NADH and (GAPDH-NADH)-LDH complex were set to 5 and to 40 sec^-1^ (Supp. Fig. 11, steps *k-4*, *k-7*, and *k9*). The NADH binding for GAPDH is eight times weaker in panel B than in panel A, what leads to accumulation of LDH-NADH, (LDH-NADH)-GAPDH, and (LDH-NAD)-GAPDH complex at the expense of (GAPDH-NADH)-LDH, LDH-GAPDH, and GAPDH-NADH complex.

(**C**) Dissociation rates for GAPDH-NADH complex (Supp. Fig. 11, steps *k-4*, *k-7*, and *k9*) have been linearly increased from 5, 10, 20, 30, 40 sec^-1^. The result is a hyperbolic increase in the ratio between the observed LDH activity and the calculated free diffusion activity. The hyperbolic increase in the ratio indicates that the increase in the off-rates affects to a different extent the channeling and the no-channeling path.

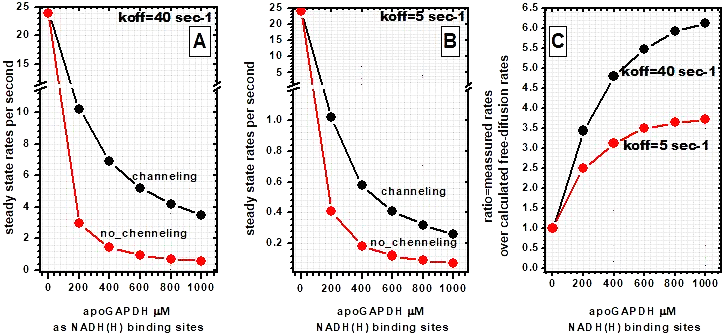

**Supp. Fig. 13 (A-C). Changes in off-rates for GAPDH-NADH complex can reproduce experimentally observed differences between rmGAPDH and byGAPDH.** The activity of 1 µM LDH with 40 µM NADH was simulated in the presence of 1, 200, 400, 600, 800, 1000 µM of GAPDH. The LDH activity in the presence of 1 µM GAPDH represents LDH reaction with free NADH because in those conditions the enzyme concentrations are too low to support the formation of LDH-GAPDH complex. The graphs show the gradual switch from diffusive to channeling reaction paths with gradual increase in GAPDH concentration and GAPDH-NADH complex (**A-B**) All rate constants are the same as in Supp. Fig. 11 A-C, except that the *off*-rate for GAPDH-NADH complex were equal to 40 sec^-1^ or 5 sec^-1^ (steps *k-4*, *k-7*, *k9* in Supp. Fig. 11A). The simulations show that higher *off*-rates for GAPDH-NADH complex can give higher rates for both the *observed LDH activity* and the *calculated free diffusion activity*. (**C**) Higher ratio between *observed LDH activity* and *calculated free-diffusion* rates can be observed with the higher *off*-rates (i.e., higher *Kd* constants) at all GAPDH concentrations. These observations are consistent with the experimentally observed differences between byGAPDH and rmGAPDH (Fig. 7-8) and indicate that the higher *off*-rates for GAPDH-NADH complex favor the channeling reaction path.

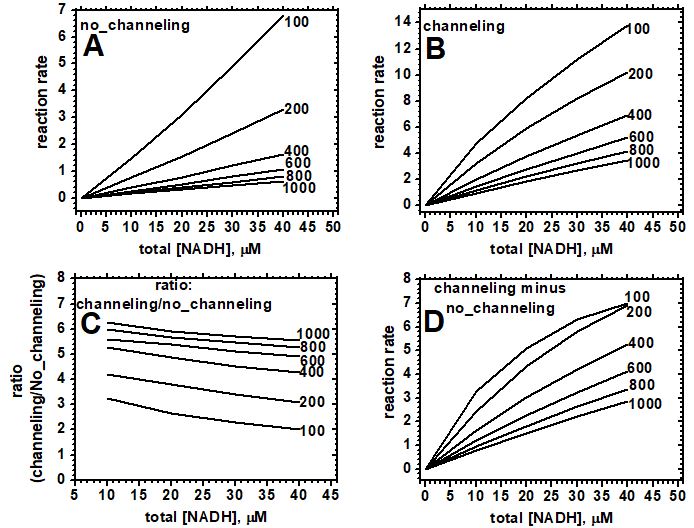

**Supp. Fig. 14 (A-D). Simulation of Michaelis-Menten curves for LDH activity with its NADH substrate for “no channeling” and “channeling” paths in the presence of different concentrations of GAPDH (the numbers next to lines).**

**(A)** The panel shows LDH activity with variable total NADH concertation and a large excess of GAPDH fixed at different concentrations (the numbers next to lines). Just as in Supp. Fig. 11A, the rate constants for steps 1 to 4 are listed in the table, while the rate constants for “the channeling steps” 5 to 12 were set to zero (i.e., no interaction between LDH and GAPDH). The simulations show that in the case of “no-channeling” apo-GAPDH acts as a competitive inhibitor of LDH activity.

(**B**) Same simulations as in panel A, except that the rate constants for “the channeling steps” 5 to 12 were set to the values listed in the table in Supp. Fig. 11 (i.e., channeling included). The figure shows that apo-GAPDH again acts as a competitive inhibitor of LDH activity; however the inhibition is now less pronounced than in “no-channeling” mechanism in panel A.

(**C**) The calculated ratios between the rates shown in panel B and panel A show that enzyme buffering experiments can detect the differences between channeling and diffusive reaction at different GAPDH and NADH concentration. The panel shows that even at extreme GAPDH and NADH concentrations there is relatively little variation in calculated R values (30% variation) which indicates that for specific enzyme pairs our ability to measure channeling can be limited by the rate constants that are specific for those enzymes, most notably *off*-rates for GAPDH-NADH complex (Supp. Fig. 12).

(**D**) Curves in panel A are subtracted from the curves in panel B to show that in the enzyme buffering experiments constant completion between channeling and no-channeling reaction path regulates the apparent Km for LDH activity in the channeling path.

### 6. Full numerical description of enzyme buffering experiments

**Supp. Table 2 A-B. NADH channeling from rmGAPDH-NADH (A) or byGAPDH-NADH (B) complex to rmLDH or phLDH (full numerical description of experiments presented in figure 7 and 8).** The presented values illustrate the design of the enzyme buffering experiments and our ability to separate the channeling reaction from the parallel reaction by free-diffusion. Free NADH concentrations were calculated using Kd constants for GAPDH values (Table 1). The calculated activity represents expected activity in the case of no channeling using equations 2-3 (methods) and Michaelis-Menten constants that depict LDH activity (Table 2).

A

| [rmGAPDH] | [NADH] | [NADH] | rmLDH activity U/mg | | | phLDH activity U/mg | | |
| --- | --- | --- | --- | --- | --- | --- | --- | --- |
| μM, site | Total μM | Free μM | Experimental | calculated | ratio | experimental | calculated | Ratio |
| 120 | 40 | 0.389 | 13.65 | 11.38 | 1.2 | 8.09 | 6.18 | 1.31 |
| 140 | 40 | 0.313 | 13.00 | 9.29 | 1.4 | 7.21 | 5.01 | 1.44 |
| 160 | 40 | 0.261 | 11.37 | 7.84 | 1.45 | 6.65 | 4.21 | 1.58 |
| 180 | 40 | 0.224 | 10.86 | 6.78 | 1.6 | 6.24 | 3.63 | 1.72 |
| 200 | 40 | 0.196 | 10.16 | 5.98 | 1.7 | 5.78 | 3.19 | 1.81 |
| 220 | 40 | 0.175 | 9.30 | 5.34 | 1.74 | 5.47 | 2.85 | 1.92 |
| 240 | 40 | 0.157 | 8.94 | 4.83 | 1.85 | 5.14 | 2.57 | 2 |
| 200 | 10 | 0.042 | 2.97 | 1.32 | 2.25 | 1.58 | 0.69 | 2.28 |
| 200 | 20 | 0.088 | 5.79 | 2.76 | 2.10 | 3.13 | 1.46 | 2.15 |
| 200 | 30 | 0.140 | 8.23 | 4.33 | 1.90 | 4.55 | 2.30 | 1.98 |
| 200 | 40 | 0.199 | 10.29 | 6.05 | 1.70 | 5.85 | 3.23 | 1.81 |
| 200 | 50 | 0.265 | 13.19 | 7.95 | 1.65 | 7.26 | 4.27 | 1.7 |

rmGAPDH-NADH complex Kd= 0.8 ± 0.06 μM.

rmLDH K_M_= 4.4 ± 0.4 μM; Vmax= 140 U/mg

phLDH K_M_= 7.8 ± 0.9 μM; Vmax= 130 U/mg

B

| [byGAPDH] | [NADH] | [NADH] | rmLDH activity U/mg | | | phLDH activity U/mg | | |
| --- | --- | --- | --- | --- | --- | --- | --- | --- |
| μM, site | Total μM | Free μM | Experimental | calculated | ratio | experimental | calculated | ratio |
| 240 | 40 | 1.6 | 113.8 | 36.7 | 3.1 | 71.6 | 21.7 | 3.3 |
| 280 | 40 | 1.3 | 109.5 | 32.2 | 3.4 | 67.5 | 18.7 | 3.6 |
| 320 | 40 | 1.1 | 109.0 | 28.7 | 3.8 | 66.0 | 16.5 | 4 |
| 360 | 40 | 1.0 | 108.6 | 25.8 | 4.2 | 64.8 | 14.7 | 4.4 |
| 400 | 40 | 0.9 | 108.2 | 23.5 | 4.6 | 62.5 | 13.3 | 4.7 |
| 440 | 40 | 0.8 | 103.6 | 21.6 | 4.8 | 63.0 | 12.1 | 5.2 |
| 480 | 40 | 0.7 | 101.7 | 19.9 | 5.1 | 60.1 | 11.1 | 5.4 |
| 480 | 10 | 0.042 | 32.55 | 5.25 | 6.2 | 18.45 | 2.80 | 6.6 |
| 480 | 20 | 0.088 | 59.83 | 10.32 | 5.8 | 34.06 | 5.58 | 6.1 |
| 480 | 30 | 0.140 | 82.13 | 15.21 | 5.4 | 47.67 | 8.36 | 5.7 |
| 480 | 40 | 0.199 | 101.68 | 19.94 | 5.1 | 60.12 | 11.13 | 5.4 |
| 480 | 50 | 0.265 | 118.86 | 24.51 | 4.85 | 72.27 | 13.90 | 5.2 |

byGAPDH-NADH complex Kd= 8.2 ± 0.5 μM.

rmLDH K_M_= 4.4 ± 0.4 μM; Vmax= 140 U/mg

phLDH K_M_= 7.8 ± 0.9 μM; Vmax= 130 U/mg

**Supp. Table 3 A-B. NADH channeling from rmLDH-NADH (A) or phLDH-NADH (B) complex to rmGAPDH or byGAPDH**. The activities measured in the assays are compared to the calculated activity which represents expected activity in the case of no channeling (equations 2-3 methods). Channeling was measured only at the highest LDH concertation to minimize the effect of allosteric regulation of GAPDH activity by the NAD+ produced during the reaction . For the same reason the apparent Michaelis-Menten constants for GAPDH activity have been calculated using NADH concentrations below 10 μM .

A

| [rmLDH] | [NADH] | [NADH] | rmGAPDH activity U/mg | | | byGAPDH activity U/mg | | |
| --- | --- | --- | --- | --- | --- | --- | --- | --- |
| μM, sites | Total μM | Free μM | Experimental | calculated | ratio | experimental | calculated | Ratio |
| 400 | 40 | 0.78 | 100.45 | 20.5 | 4.9 | 69.0 | 11.7 | 5.9 |

rmLDH-NADH complex Kd= 7.2 ± 0.07 μM;

rmGAPDH K_M_= 3.4 μM ± 0.8; Vmax= 110 U/mg

byGAPDH K_M_= 6.2 μM ± 0.7; Vmax= 105 U/mg

B

| [phLDH] | [NADH] | [NADH] | rmGAPDH activity U/mg | | | byGAPDH activity U/mg | | |
| --- | --- | --- | --- | --- | --- | --- | --- | --- |
| μM, site | Total μM | Free μM | Experimental | calculated | ratio | experimental | calculated | ratio |
| 400 | 40 | 0.46 | 45.9 | 13.1 | 3.5 | 31.2 | 7.25 | 4.3 |

phLDH-NADH complex Kd= 4.2 ± 0.09 μM;

rmGAPDH K_M_= 3.4 μM ± 0.8; Vmax= 110 U/mg

byGAPDH K_M_= 6.2 μM ± 0.7; Vmax= 105 U/mg

### 7. Materials and Methods

*Materials*. NAD^+^ and NADH were of 99% or better purity as listed. D(-)3-phosphoglyceric acid (98% pure as listed) was obtained as the tri(cyclohexylammonium) salt. ATP was the disodium salt grade I. BSA was RIA grade, fraction V powder. Activated charcoal was hydrochloric acid-washed, cell culture tested. Ammonium sulfate suspensions of phLDH, pmLDH, were purchased from Sigma Chem. Co. or MP Biomedicals (Costa Mesa, CA). byGAPDH and rmGAPDH were obtained as 50% glycerol or ammonium sulfate suspensions from Sigma Chem. Co., or prepared in-house . Polyclonal goat anti-rabbit LDH antibodies were purchased from ICN Pharmaceuticals. NADH stock solutions were prepared in 120 mM Na_2_CO_3_ in a light isolated container and used within a week of preparation. NAD^+^ was prepared fresh before each experiment. All molecular dynamics calculations were carried-out on Bullx DLC 720 supercomputer at the Center for Advanced Computing and Modeling at the University of Rijeka (http://cnrm.uniri.hr/). A typical calculation used 50 nodes that were integrated by Infiniband FDR 54 Gbps connections. Each node has two Intel Xeon E5-2690 v3 processors, NVIDIA K40 GPU and 64GB RAM.

*Apo-enzyme preparations*. (NH_4_)_2_SO_4_ suspensions of purified rmGAPDH and byGAPDH have been prepared in our laboratory using established protocols or purchased from Sigma Chem. Co. Specific activity of purified enzyme was measured in direction of NADH oxidation. The assay mixture had 3 mM 3‑phosphoglycrate, 1 mM ATP, 2 mM MgCl_2_, 10 U/ml of 3–phosphoglycerate kinase, and 100 uM NADH . Specific activity of purified byGAPDH was 95-110 U/mg while rmGAPDH had 100-120 U/mg for rmGAPDH. An enzyme unit (U) is defined herein as the amount of enzyme producing or using 1 μmol of NADH/min in the specified assay conditions .

Prior to each experiment, the enzyme solutions were centrifuged to remove excess of (NH4)2SO4, and filtered in the assay buffer through a G-50 column. Following the gel filtration, byGAPDH had absorbance ratios at 280/260 nm equal to 2.00 ± 0.02 which indicates that the enzyme was NAD(H) free . rmGAPDH showed an absorbance ratio of 1.81 ± 0.01 which indicates that the enzyme still had some tightly bound NAD(H) that could not be removed with gel filtration . Subsequent treatment with acid washed charcoal can remove the tightly bound NAD(H) and increase the ratio to 1.98 ± 0.01 . Complete removal of NADH molecules leads to dissociation of rmGAPDH tetramers that we could detect with AUC and results in scattering at high GAPDH concentration in our enzyme buffering test . Thus we did all our experiments with rmGAPDH with 280/260 nm equal to 1.8, which indicates that the enzyme still had tightly bound NADH on two of its four subunits . The tightly bound NADH does not participate in the reaction in enzyme buffering tests since there is no LDH activity in 30-60 seconds of assay time when GAPDH and LDH are mixed together in the absence of added NADH.

(NH_4_)_2_SO_4_ suspensions of phLDH and pmLDH were purchased from Sigma and prepared using gel filtration just as rmGAPDH and byGAPDH. For all purified proteins concentrations were calculated by measuring absorbance at 280 nm in 1mm wide cell using following absorptivity (M^-1^cm^-1^): phLDH 1.7x10^5^; rmLDH 1.75x10^5^; rmGAPDH 1.16x10^5^; byGAPDH 1.3x10^5^. The molar absorptivity, pI, and molecular mass (Mr) values were calculated (www.expasy.ch) from the available sequences (http://www.uniprot.org/).

*Molecular dynamics calculations*. All-atom molecular dynamics calculations used the GROMACS 5.1.4 program package as we have previously described . Briefly, NAD(H) molecules from PDB files were processed using ACPYPE10, an interface for Antechamber (part of AmberTools11) that can generate topology types for GAFF force fields . Protein PDB coordinates were processed with pdb2gmx using Amber99SB force field . A cubic solvent box (30 nanometers) was used with TIP3P model for water molecules plus 150 mM NaCl and additional ions that were required for neutralization. The prepared system was minimized using a combination of steepest descent and conjugate gradient algorithms. When the most stable state was achieved the temperature was introduced and the system was equilibrated to 310 K (NVT equilibration, tcouple V-rescale, columbtype PME long-range, cutoff-scheme Verlet, rcoulomb and rvdw set to 1.0). The pressure was equilibrated to 1 atm (NPT equilibration, pcoupl Parrinello-Rahman, tcouple V-rescale, cutoff-scheme Verlet, columbtype PME long-range). No restraints were used for the protein or the ligand when the system was minimized, but in NPT and NVT equilibration protein and ligand were mutualy restrained to prevent the integrity of the complex. The outputs of minimization and equilibration MD simulations provide insight into the potential energy of the system based on minimization of the temperature in NVT simulation, and minimization of the pressure in NPT simulation.

Typical simulations had about 1.5 million atoms, 50-150 million steps with time step set to 2 femtoseconds Total duration was 100 to 300 nanoseconds with the structures sampled for visualization every 0.1 nanosecond. Large simulation boxes (30 nanometers) were used to avoid attractive or repulsive forces created by the periodic boundary boxes. The large boxes can also provide enough space for the two tetramers to dissociate apart (supplement movie 1). Different initial simulation set-ups were used to explore the validity of simulations. For example, the simulations were repeated with active site loop on LDH in open and closed position, with different initial distances between the interacting proteins, and with different rotations between the LDH and GAPDH tetramers in the interaction plane. Following the simulations, the number of binding interactions was calculated using built-in GROMAC functions. The simulations that showed the highest number of binding interactions have been repeated multiple times to show variability in the number of binding interactions.

*Adaptive Poisson-Boltzmann Solver (APBS) calculations.* All electric fields maps were calculated using Adaptive Poisson-Boltzmann Solver (APBS) approach . Protein coordinates were converted from PDB format to PQR format using PDB2PQR application and PEOEPB force filed with PROPKA set at pH=7.2. NAD(H) molecules from PDB files were protonated using GAFF fields. Potential maps were calculated in aqueous 150 mM NaCl solutions using single Debye-Hückel boundary conditions.

*Rigid body protein-protein docking*. GAPDH and LDH docking models were prepared using the DOT version 2.0.1 suite and allied tools installed on NREL’s Peregrine system . Starting from a reference snapshot from all-atom molecular dynamics simulations (Fig 1), structures were preprocessed as required to create the stationary GAPDH model. The mobile LDH model sampled through rotational coordinates in X degree increments around the GAPDH center (to sample center-of-mass orientation of the two proteins relative to one another) and its center (to sample relative binding surface interactions between the two proteins).

The DOT2 package uses as its base metric electrostatic and van der Waals energies parameterized similarly to all-atom force fields like AMBER . To process the protein input structures through the DOT2 preprocessing pipeline without errors, it was necessary to parameterize the GAPDH-bound NADH molecule for electrostatics, van der Waals forces, and desolvation energies in the ACE and ODA models. Electrostatics and van der Waals parameters were derived from the Parm14 IPolQ van der Waals parameters distributed with AMBER 18 , with charges published online by Ulf Ryde. Heavy atom type mappings (both ACE and ODA do not parameterize hydrogens) among AMBER, ACE, and ODA are shown in Table Supp 1.

Docking runs were performed using DOT2’s finest rotational mesh with samples 4° apart, and the default 1 Å translational grid spacing. Post-processing focused on the 2000 lowest energy structures found as judged by electrostatics and van der Waals energies alone. The associated poses were then rescored with potentials including either the ACE or ODA desolvation models.

The VMD program package version 1.9.3 was used to visualize protein structures. LDH poses were encoded by arrows within each chain (Fig Supp 1), extending from an atom inside the substrate binding site to a loop atom on the protein surface approximately at the exit of the binding site. In this way, the arrows roughly encode the paths one would expect NADH to take in the process of binding to LDH, and each LDH pose is represented by two arrows, one for each polypeptide chain.

The lowest energy 100 structures from each energetic model were then analyzed, to assess any unphysical positional biases arising from the energetic models, and any potential spatial clustering with respect to low-energy LDH poses. In addition, the base 2000 structures were filtered based on the dual distance constraints of:

1. ≤ 16 Å between GAPDH Arg77 CZ and LDH Phe 188 CG, and

2. ≤ 4 Å between GAPDH Glu76 CD and LDH Arg 111 NH2

Together, these constraints were intended to select for poses comparable to the initial reference structure. These were applied to chains C (LDH) and P (GAPDH) only; however, as homomultimers, one would naturally expect symmetric results associated with chains B and Q, respectively. Analysis of relative energetics was done through Python scripting within a Jupyter notebook.

*Coarse grained molecular dynamics studies*. Coarse-grained molecular dynamics simulations CG MD simulations were was generated using CHARMM-GUI Martini Solution Builder with MARTINI 2.2 force field . Energy minimization was carried out with 6000 cycles of steepest descent. The systems were relaxed using 2 equilibration steps of 2 and 10 ns. Production simulations were performed for 10 µs with 20 fs time step. The temperature was set to 315.15 K using V-rescale coupling, and the pressure was set to 1.0 bar using semi-isotropic Berendsen coupling. The temperature was set to 310 K using V-rescale coupling, the pressure was set to 1.0 bar using semi-isotropic Berendsen coupling, pcoupl set to Parrinello-Rahman, columbtype set to Reaction-Field. A cubic solvent box (26.5 nanometers) was used with TIP3P model for water molecules plus 150 mM NaCl and additional ions for neutralization. The systems were relaxed using equilibration steps and simulations had 500 million steps, with time step set to 20 fs, in total duration of 10 µs.

*Fluorescence measurements of NADH binding affinity for rmGAPDH and byGAPDH*. NADH binding to GAPDH molecules was measured using four different types of fluorescence measurements to achieve maximal accuracy . Protein fluorescence measurements had excitation 290nm and emission at 335nm. NADH fluorescence had excitation at 340 nm and emission at 460 nm. Protein-NADH FRET measurements had extraction 290 nm and emission at 460 nm. NADH anisotropy had excitation at 340nm and emission at 460 nm. All fluorescence measurements were measured in a microcuvette in a total volume of 80 μl of assay buffer at 25° C using PE LS-50B fluorescence instrument. In all measurements dissociation constant (Kd) for GAPDH-NADH complex was calculated using nonlinear regression. For GAPDH or NADH fluorescence quenching experiments that measured a decrease in the signal from free GAPDH or free NADH we used following binding equation:

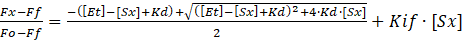
eqn. 1a

In FRET and fluorescence anisotropy measurements which measure signal from GAPDH-NADH complex we used equation:

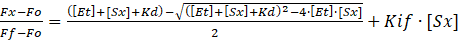
eqn. 1b

Where [*Et*] represents total GAPDH concentration in terms of NAD(H) binding sites, while [*Sx*] represents NADH at concentration *x*. *Fo* represents initial fluorescence, *Fx* measured fluorescence at NADH concentration [*Sx*], *Ff* represent final fluorescence at full saturation with NADH. Finally, *Kif* represents the slope of decrease in fluorescence caused by the inner-filter effect that is due to the tail end of the NADH-adenine absorbance peak. The equations 1 a-b were used in nonlinear regression with *Ff, Kd*, and *Kif* as free fit parameters, while *Et, Fo, Fx* and [*Sx*] represent measured values specific for each experimental set-up.

The slope of *Kif* constant that was calculated by nonlinear regression as a free fit parameter was compared to four different control measurements. In first control, the GAPDH solution used for fluorescence measurements was also subjected to uv-vis spectrophotometry to measure the increase in absorbance at the excitation wavelength caused by the addition of NADH. In the second control, GAPDH was replaced in the fluorescence measurements with a solution of free tryptophan in concentration that gives identical initial fluorescence *Fo* as the GAPDH. In third control, GAPDH was replaced with BSA solution with identical initial fluorescence *Fo*. In the fourth control, NADH binding to GAPDH was measured at three different GAPDH concentrations, because our ability to calculate *Kif* from *Fi* and *Ff* measurements depends on the GAPDH concentration used in the measurements. If the *Kif* was calculated correctly the *Kd* values from different fluorescence measurements are to within experimental error identical. The maximal experimental variability that was tolerated in our *Kd* measurements was ± 10%.

The measured binding constants for rabbit and baker’s yeast GAPDH are within a factor of 3 equal to the previously reported values and more accurate since we used multiple methods in the measurements and the data analysis (Table 1 vs. ).

*Enzyme buffering measurements*. LDH activity was measured by following NADH oxidation in the presence of 630 μM pyruvate which results in a decrease in NADH absorbance at 340 nm. The changes in absorbance were measured with Shimadzu UV-VIS 160 Spectrophotometer, HP 8452A Diode Array Spectrophotometer or UV-2700 UV-Vis double beam spectrophotometer. The molar absorptivity for NADH is 6.22x10^3^ M^-1^cm^-1^. In all activity measurements the experimental conditions have been adjusted to achieve precession greater than 2.4%.

The assay buffer was 50 mM Tris/HCl pH=7.4, 2 mM EDTA-Na, 1 mM DTT, and 0.5 mg/ml BSA. A low ionic strength buffer was chosen to favor detection of electrostatic channeling . The assay mix was prepared in a microcuvette in a total volume of 80 μL and its temperature was equilibrated in the cell holder to 25° C. During the incubation the absorbance at 340 nm was monitored to confirm the baseline stability. NADH was added next and the absorbance was monitored to confirm that GAPDH solution does not have any intrinsic NADH oxidize activity. If the base line was stabile 630 μM of pyruvate was added next. The resulting absorbance was measured to confirm that there is no contaminating LDH activity in the GAPDH solution. If the baseline was stabile the reaction was started by adding between 2 to 20 nM of LDH depending on the desired experimental sensitivity.

In assays with NADH concentration between 40 to 50 μM the steady-state rates were measured by following the initial linear decrease in absorbance. In the assays with NADH concentration below 30 μM two different approaches were used to achieve desired reproducibility. With the high sensitivity spectrophotometer we were able to capture the initial linear steady state even at the lowest NADH concentrations at 10 μM. With low sensitivity spectrophotometers the rates were calculated using exponential equations to calculate the rate of decrease in NADH absorbance . byGAPDH solutions were stable even at the highest enzyme concentration tested at 16mg/ml. rmGAPDH gradually precipitates at the level above 8.4 mg/ml, which leads to scattering and detectable increase in absorbance. Thus, the LDH concentrations were adjusted to maximize the ratio between the changes in absorbance caused by NADH oxidation and by scattering. The scattering was than subtracted as a minor component in the measured changes in NADH absorbance.

The free NADH concentration [NADH]f in the assay mix can be calculated using Kd for GAPDH:

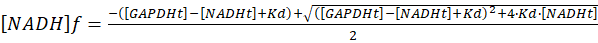
eqn. 2

Where [GAPDH] concentrations represent concentration in terms of NADH binding sites, while [GAPDH-NADH] represents the concentration of the complex, based on the equation [GAPDH] + [GAPDH-NADH] = [GAPDH]total. *Kd* is the GAPDH-NADH dissociation constant calculated from the fluorescence measurements. The calculated [NADH]free was used to calculate expected LDH activity in case of no channeling between LDH and GAPDH:

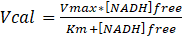
eqn. 3

Vmax and Km are the values for Michaelis-Menten constants for NADH oxidation reaction with LDH. All LDH assay mixtures were prepared with 630 μM pyruvate. Specific activities of phLDH, rmLDH, were 130 ± 15 and 430 ± 30 U/mg, respectively.

*Sedimentation Velocity AUC Experiments.* We used Beckman XL-A instrument, An-60-Ti rotor, and three sample cells with quartz windows and charcoal-epon centerpieces with two sectors 12 mm optical path length. For each cell, 340 μL enzyme samples (6.0 µM) were loaded in one sector, and 350 μL buffer in another. Two cells had individual enzymes, and the third cell had the enzyme mixture at the same loading concentration. The experiments started with the prescans at 3000xg, and the sum of absorbances of the individual enzymes was compared to the absorbance of the mixture. The sedimentation profiles were measured for 7 hours at 40,000xg. Single scans with radial increments of 30 μm have been recorded every 240 sec. SEDNTERP was used with protein sequences to calculate molecular mass, partial specific volume, solvent densities, and viscosities .
